# A Broad-Spectrum Chemokine Inhibitor Prevents Preterm Labor in Mice by Suppressing Inflammation Induced by Intra-Amniotic Injection of Interleukin-1 alpha

**DOI:** 10.64898/2026.03.04.709657

**Authors:** Adam Boros-Rausch, Nicole Ballan, Isabella Celik, Anna Dorogin, Zoe E Gillespie, Jennifer A Mitchell, David Grainger, David Fox, Oksana Shynlova, Stephen Lye

**Author notes:** Address for Correspondence: Dr. Oksana Shynlova, Lunenfeld-Tanenbaum Research Institute at Mount Sinai Hospital, 25 Orde Street, Suite 6-1017 Toronto, Ontario, Canada M5G 1X5. Equal contributions (Oksana Shynlova, Stephen Lye).

## Abstract

Preterm birth (PTB) is a leading cause of perinatal and infant mortality worldwide. PTB can be induced by systemic maternal or intra-uterine infection or by sterile intra-amniotic inflammation driven by alarmins such as interleukin-1α (IL-1α). We reported earlier that a Broad-Spectrum Chemokine Inhibitor (BSCI) prevented PTB in murine and non-human primate models of infection-mediated preterm labor. Here, we investigated whether BSCI can prevent PTB in pregnant C57BL/6 mice following ultrasound-guided intra-amniotic injection of IL-1α (400 ng per sac) on gestational day (GD)16.5. Half the mice received BSCI (10 mg/kg, intravenous daily) beginning GD15.5 and through to term. The impact of IL-1α alone or IL-1α plus BSCI was assessed on (i) injection-to-delivery interval, and fetal survival; (ii) cytokine and chemokine levels in maternal plasma and amniotic fluid and inflammatory gene expression in maternal and fetal tissues (Real-Time RT-qPCR); (iii) global transcriptomic profiling of myometrial tissues at 2 and 24 hrs (RNA sequencing) together with examination of myometrial chromatin accessibility (Assay for transposase accessible chromatin sequencing) at 24 hrs; (iv) uterine leukocyte infiltration (immunofluorescence). Pretreatment with BSCI i) reduced the incidence of IL-1α-induced PTB; (ii) significantly attenuated cytokine and chemokine signals in maternal plasma, myometrium, decidua, and placenta, and amniotic fluid; (iii) suppressed myometrial contraction-associated genes, including *Nfkb1, Ptgs2, Akr1c18*, and *Gja1*; (iv) prevented global IL-1α-induced changes in myometrial gene expression and chromatin accessibility (v) reduced uterine macrophage (F4/80+) counts and prevented the increase in pro-inflammatory M1-like macrophages observed with IL-1α-treatment. Taken together, BSCI reduced the occurrence of PTB in mice, maintained uterine quiescence by suppressing intra-amniotic inflammation and genome-wide changes in labor gene expression and chromatin accessibility. BSCI represents a promising therapeutic approach for PTB prevention in high-risk pregnant women.

## INTRODUCTION

Preterm birth (PTB), defined as delivery before 37 weeks of gestation, is the leading cause of neonatal morbidity and mortality worldwide, accounting for nearly 15 million births annually and contributing to more than one million infant deaths each year (1, 2). Survivors of PTB often face significant long-term health complications, including respiratory distress syndrome, bronchopulmonary dysplasia, necrotizing enterocolitis, cerebral palsy, and cognitive or sensory impairments (3, 4). Despite decades of research, the global rate of PTB remains unchanged, representing an urgent and unmet need in maternal-fetal medicine. PTB is a multifactorial syndrome with diverse etiologies, including infection, uteroplacental ischemia, cervical insufficiency, and immune dysregulation (5). While microbial invasion of the amniotic cavity is a well-documented cause of infection-induced PTB, a significant subset of women with preterm labor exhibit elevated inflammatory markers in amniotic fluid despite negative microbiological cultures (6). This clinical condition, termed “sterile” intra-amniotic inflammation (sterile IAI), has been observed in approximately one-third of women with spontaneous PTB (sPTB) and intact membranes (6–10). Among sPTB, IAI is a major precipitating factor (11–13).

Sterile IAI is believed to be initiated by endogenous molecules called damage-associated molecular patterns (DAMPs), or alarmins, that are released during cellular stress or necrosis (14–16). These include interleukin (IL)-1α, high-mobility group box 1 (HMGB1), S100B, and heat shock proteins, which activate innate immune responses through Toll-like and NOD-like receptors (13, 16, 17). Among these, IL-1α has been recognized as an essential cytokine in the pathogenesis of sterile IAI (8, 16, 18–22). It is constitutively expressed in many cell types and functions both as a transcriptional regulator and as an extracellular alarmin upon release from necrotic cells. Elevated levels of IL-1α in amniotic fluid have been correlated with the onset of both term labor and sPTB and are linked to adverse neonatal outcomes (23–26). Experimental mouse models have demonstrated that intra-uterine and intra-amniotic administration of IL-1α induces inflammatory activation of fetal membranes, preterm labor contractions, and fetal brain injury (16, 27, 28).

Current clinical management of PTB remains limited, primarily focusing on delaying delivery by tocolytics while using corticosteroids to enhance fetal lung maturation. However, these interventions do not address the underlying immunopathology. Given the central role of inflammation in both infectious and sterile PTB, targeting pro-inflammatory pathways represents a promising strategy for therapeutic measures. Chemokines, a family of small cytokines, are essential regulators of leukocyte trafficking and activation, and are strongly upregulated during intrauterine inflammation (29, 30). While specific chemokine receptor antagonists have been trialed in models of cancer, autoimmune disease, and HIV, their narrow specificity often limits efficacy in complex inflammatory networks (31–37). To overcome this limitation, Broad-Spectrum Chemokine Inhibitors (BSCIs) have been developed to simultaneously target multiple chemokine signaling pathways without generally suppressing host immunity (38). BSCIs have demonstrated efficacy in mitigating inflammatory conditions, including atherosclerosis, HIV replication, surgical adhesion formation, lung disease, ischemia, rheumatoid arthritis, endometriosis, and psoriasis (33, 39–45). Importantly, we and others have demonstrated that BSCI administration effectively prevents infection-induced PTB in both murine and non-human primate models, significantly reducing cytokine production, leukocyte infiltration, and uterine contractility without compromising fetal viability (46, 47).

Our earlier studies demonstrated significant anti-inflammatory effects of BSCI relevant to PTB mechanisms. The BSCI compound FX125L inhibited the migration of human primary leukocytes *in vitro,* (including neutrophils, monocytes, and lymphocytes) across endothelial cell layers, indicating their potential to limit immune cell infiltration into uterine tissues *in vivo* (48, 49). Our mechanistic investigations further revealed that BSCI suppressed key intracellular signaling events in human uterine smooth muscle cells (myocytes), including NF-κB activation and secretion of chemokines IL-8 and CCL2 (50). This disruption impaired macrophage–myocyte communication and reduced *in vitro* myocyte contractility, supporting the potential of BSCIs to modulate inflammatory cascades that contribute to labor onset. BSCI also alters the secretome of human myocytes *in vitro,* promoting the polarization of human macrophages toward the anti-inflammatory M2 phenotype (51). M2 macrophages, characterized by high expression of CD206 and TGF-β, plays a crucial role in maintaining tissue homeostasis and suppressing inflammation (52–54). In contrast, M1 macrophages express IL-1β, TNF-α, and generate reactive oxygen species (ROS), which are known to promote labor onset and fetal injury (52, 55–57). Studies in human and murine tissues have demonstrated that M2 macrophages are predominant at the maternal-fetal interface during late gestation but decline with the onset of labor, while M1 polarization increases in both term and preterm labor (58–60). Macrophage depletion or the adoptive transfer of M2 macrophages has been shown to delay preterm labor in rodent models, reinforcing their functional importance in preserving uterine quiescence (60–66).

Given this context, we hypothesized that the BSCI compound FX125L could prevent IL-1α-induced sterile IAI and sPTB in mice by suppressing inflammatory signaling and preserving M2 macrophage polarization *in vivo*. In this study, we employed a well-characterized mouse model of sPTB induced by ultrasound-guided intra-amniotic injection of alarmin IL-1α (16). We tested the efficacy of BSCI administered systemically in preventing PTB. We evaluated (i) the IL-1α injection-to-delivery interval, and fetal viability,; (ii) cytokine and chemokine expression in maternal tissues and amniotic fluid; (iii) changes in the myometrial transcriptome and genome regulatory landscape; and (iv) monocyte infiltration and macrophage phenotypes within the uterus. Our findings demonstrate that BSCI reduced reproductive tissue inflammation in pregnant mice, thus preserving M2 macrophage dominance in the murine uterus. These results support the therapeutic potential of BSCI for preventing IAI-driven PTB in high-risk pregnant women.

## MATERIALS AND METHODS

### Ethics Approval

All experimental procedures were approved by the Animal Care Committee of The Centre for Phenogenomics (TCP, Toronto, Canada) (AUP #0164H) and conducted in accordance with the guidelines of the Canadian Council on Animal Care. Animals were housed in a pathogen-free, humidity-controlled 12 hr light, 12 hr dark cycle in the TCP animal facility with free access to food and water. Guidelines set by the Canadian Council for Animal Care were strictly followed in handling mice. Young (8–12 weeks old) virgin female C57BL/6 inbred mice were mated with males; the morning of vaginal plug detection was designated as gestational day (GD)1. The ratio of male to female was 1:1. The gestational length in C57BL/6 wild-type mice is 19.5 days on average (range 19.1–19.9), and term delivery under these conditions occurred during the evening of GD19 or the morning of GD20.

### Experimental Design

#### BSCI Administration

Pregnant C57BL/6 mice (n = 100) were randomly assigned to receive either BSCI or saline (vehicle) through intravenous tail vein injection. The drug was first administered to pregnant mice on GD15.5 at a dose of 10 mg/kg in 100 μl of saline, used in our previous animal studies (46, 47), and continued daily until delivery. On GD16.5, mice received intra-amniotic IL-1α or saline infusions (**Fig. 1**) to induce PTB (as described below).

**Figure 1.**
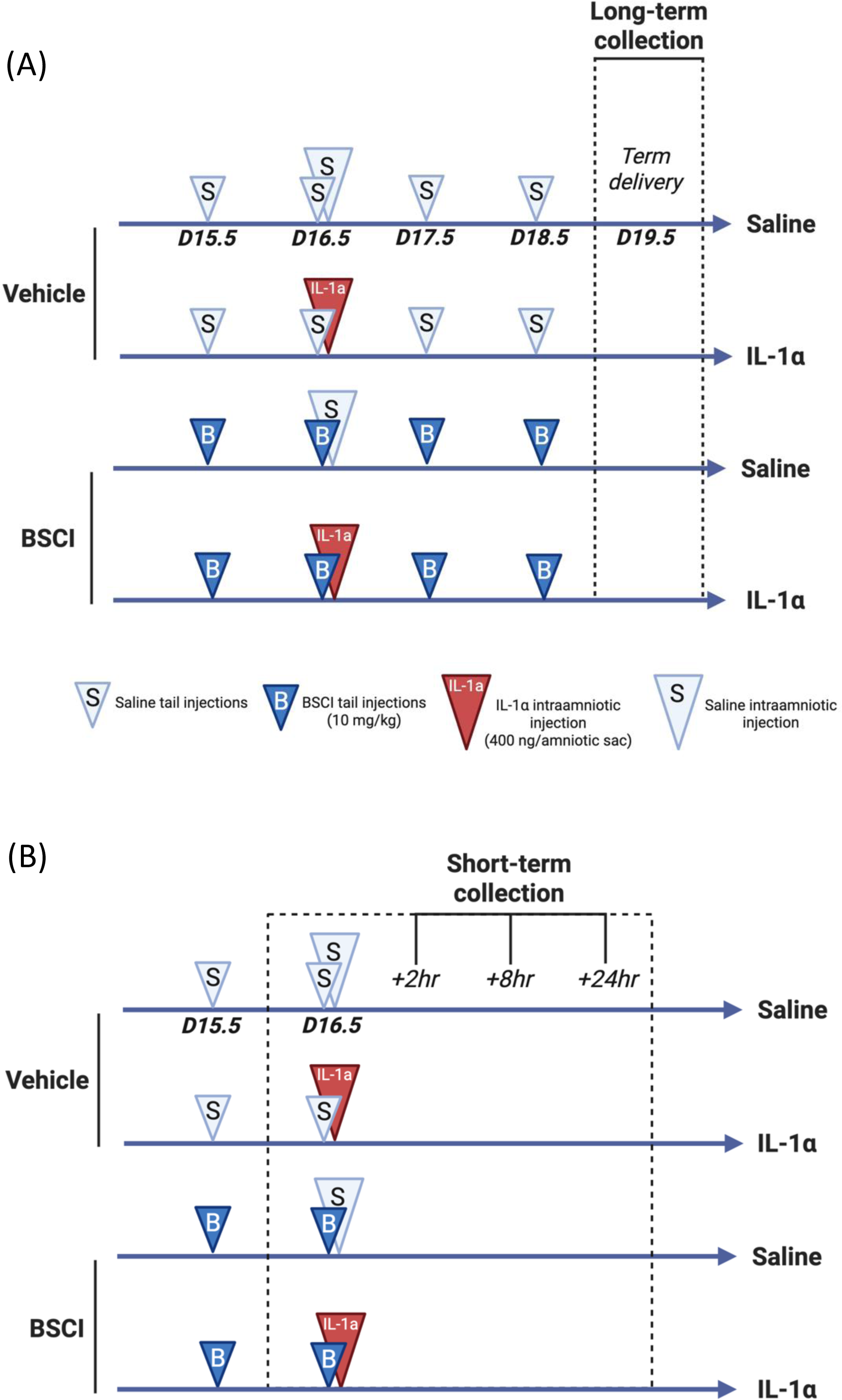
Scheme of injections and tissue collection. On GD15.5, pregnant mice [‘Vehicle’ group (n = 50) and ‘Broad Spectrum Chemokine Inhibitor’ (BSCI) group (n = 50)] were injected intravenously with the first dose of BSCI (10 mg/kg/day) or vehicle (saline). In 24 hrs, on GD16.5, pregnant mice received a second injection of BSCI or vehicle. At the same time, half of the vehicle group (n = 25) and half of the BSCI group (n = 25) received ultrasound-guided intra-amniotic infusions of IL-1α (IL-1α; 400 ng/amniotic sac). The second half of the vehicle group (n = 25) and half of the BSCI group (n = 25) received an infusion of sterile saline. The IL-1α injections were administered into each amniotic sac (number of fetuses was 5-9). **(A)** <u>Long-term effect of BSCI.</u> Pregnant mice were observed until delivery to record the PTB rate. All BSCI-treated animals from the IL-1α and saline groups that did not deliver preterm were given daily injections of BSCI. Mice from all four study groups that carried the pregnancy to term were killed after delivery on GD19.5. The number of live pups per litter, birth weights, and placental weights were recorded. **(B)** <u>Short-term effect of BSCI.</u> In a replicate group of animals, pregnant BSCI-treated and vehicle-treated mice were sacrificed 2,8 and 24 hr post-IL-1α induction. On GD16.5, two hrs after IL-1α injection (t = 2 hr), the first six animals from each groups were killed, and maternal and fetal tissues were collected for analysis. After 8 hrs (t = 8 hr), six animals from each group were killed and maternal and fetal tissues were collected for analysis. The remaining animals (n=6/group) were killed during PTB, or exactly 24 hrs after the administration of IL-1α (t = 24 hr), and tissues were collected for biochemical and immunohistochemical evaluation.

#### Ultrasound-guided Intra-amniotic Administration of IL-1α

Pregnant wild-type mice were anesthetized on GD 16.5 by inhalation of 2–3% isoflurane (Aerrane, Baxter Healthcare Corporation, Deerfield, IL, USA) and 1–2 l/min of oxygen in an induction chamber. Anesthesia was maintained with a mixture of 1.75–2% isoflurane and 2.0 l/min of oxygen during the ultrasound procedure, which was performed using the Vevo® 2100 Imaging System (VisualSonics Inc., Toronto, Ontario, Canada). Mice were positioned on a heating pad and stabilized with adhesive tape. Fur removal from the abdomen was accomplished by applying Nair depilatory cream (Church & Dwight Co., Inc., Ewing, NJ, USA) to that area. Respiratory and heart rates were monitored by electrodes embedded in the heating pad. An ultrasound probe was anchored and mobilized with a mechanical holder, and the transducer was slowly moved toward the Aquasonic CLEAR ultrasound gel (Parker Laboratories, Inc., Fairfield, NJ, USA) applied on the abdomen. According to previous optimized conditions by Motomura et al. (16), ultrasound-guided intra-amniotic injection of IL-1α (Cat #200-LA, R&D Systems, Inc., Minneapolis, MN, USA) at concentrations of 400 ng per 25 µl of sterile 1× PBS (Fisher Scientific Bioreagents, Fair Lawn, NJ, USA) was performed in each amniotic sac using a 30-gauge needle (BD PrecisionGlide Needle, Becton Dickinson, Franklin Lakes, NJ, USA). The syringe was stabilized with a mechanical holder (VisualSonics). Control wild-type dams were injected with 25 µl of saline. The number of animals injected per group is shown in each figure legend.

#### Birth Outcomes

All animals that carried pregnancy to term (i.e., GD18.5-19.5) were killed during or after TL: n = 6 in saline group, n = 9 in BSCI group, n = 9 in IL-1α/BSCI group (**Fig. 1A**). Our criteria for labor were the delivery of at least one pup. Mice that delivered preterm (n = 8 in the IL-1α group and n = 5 in the IL-1α/BSCI group) were killed by carbon dioxide inhalation during PTL. The uterus of each female mouse was removed, opened longitudinally and the total number of fetuses, their vital signs, and fetal and placental weights were accounted.

#### Pregnancy Outcomes

To evaluate the short-term effect of BSCI on cytokine expression, we collected maternal and fetal tissues at predetermined times (2, 8, and 24 hrs after the IL-1α injection, n = 5–10/group (**Fig. 1B**). (1) Maternal blood was obtained by aortic puncture in a lithium-heparin microtainer (Microvette, Sarstedt, Germany). Plasma was isolated by centrifugation for 5 min. at 2000 × g, and the upper phase was collected and frozen in liquid nitrogen until assayed. (2) Maternal liver was collected. (3) The uterus was placed into ice-cold PBS, bisected longitudinally, and dissected away from both pups and placentas. Decidua basalis was cut away from the myometrial tissue and pooled from all implantation sites. (4) Myometrium from both uterine horns was collected and pooled. The decidua parietalis was carefully removed from the myometrial tissue by mechanical scraping on ice. Fetal tissues: (5) Amniotic fluid was collected from all gestational sacs, centrifuged for 10 min. at 5000 × g, the supernatant was transfrerred to another tube; (6) three placentas were randomly pooled from both uterine horns. All mouse tissues were flash-frozen in liquid nitrogen and stored at −80°C. Twenty-four hrs after IL-1α or vehicle administration, part of the uterus (n = 4/group) was collected for immunohistochemistry: one intact uterine horn was cut into 10–12 mm segments and placed in Optimal Cutting Temperature compound (OCT) on dry ice. Samples were stored at −80°C until sliced on the cryotome. Sections of 5 μm thickness were collected on superfrost plus slides (Fisher Scientific, Nepean, ON, Canada).

#### Reverse Transcriptase Quantitative Polymerase Chain Reaction (RT-qPCR)

Total RNA was extracted from the frozen mouse liver, myometrium, decidua and placentas using TRIZOL (Gibco BRL, Burlington, ON, Canada) according to the manufacturer’s instructions (n = 6/group). RNA samples were column-purified using RNeasy Mini Kit (Qiagen, Mississauga, ON, Canada) and treated with DNase I (Qiagen) to remove genomic DNA contamination. The process was quality-controlled by measuring yield (μg), concentration (μg/μl), and A260:280 ratios via spectrometry using Nanodrop ND-1000. RNA sample integrity was assessed using the Experion system (Bio-Rad, Mississauga, ON, Canada); all samples had RIN > 8. cDNA synthesis was performed per manufacturer’s protocol (iScript cDNA synthesis kit; Bio-Rad). RT-qPCR was performed with Luminoct SYBR Green QPCR READYMIX (Sigma-Aldrich), CFX-384 Real Time System C1000 Thermal Cycler (Bio-Rad) and specific pairs of primers (see **Table 2**). Aliquots (5 ng) of cDNA were used for each PCR reaction run in triplicate. A cycle threshold (Ct) value was recorded for each sample. Each gene was normalized to the expression of four housekeeping genes (*Ppia, Tbp, Hprt, Gapdh*) by CFX Manager software (version 2.1). Relative gene expression for IL-1α-treated animals was presented as the average fold change relative to the vehicle sample collected 2 hrs after saline administration, using the comparative Ct method.

#### Luminex Assay

To assess protein concentration of both myometrium and decidua (n = 6/group), frozen tissue samples were crushed under liquid nitrogen and homogenized in Bicine lysis buffer (25 mM Bicine, 150 mM NaCl, pH 7.6) with protease and a phosphatase inhibitor cocktail (Thermo Fisher Scientific Inc., Waltham, MA, USA). Samples were spun at 12,000 g for 15 min. at 4°C, the supernatant was transferred to a fresh tube to obtain a crude protein lysate and stored at −20°C until assayed. Total protein concentrations were determined using the Bio-Rad assay (Bio-Rad). Two hundred and fifty micrograms of protein from tissue homogenates of each myometrial sample were used for the multiplex assay. Tissue cytokine levels were quantified using Bio-plex Pro mouse Cytokine 23-plex array kit (Cat# M60009R DPD) (Bio-Rad). Plasma and amniotic fluid cytokine levels (n = 5–6/group) were quantified using Mouse Luminex Discovery 10-plex Assay (Bio-Techne, Cat# LXSAMSM). All multiplex assays were performed on the Luminex200 system and Bioplex HTF (Bio-Rad). Standards and each sample were analysed in duplicate. Data analysis was performed using Bio-Plex Manager, version 5.0 (Bio-Rad) and presented as concentrations (pg/ml).

#### Immunofluorescence Staining

Firstly, frozen tissue sections were fixed using 4% paraformaldehyde (PFA) and washed three times in PBS without Ca^2+^ and Mg^2+^ (PBS-/-), followed by serum-free protein block (DAKO) to prevent non-specific antibody binding. Then, the primary antibodies were applied and incubated overnight at 4. F4/80 rat anti-mouse Abs (Cat# MCA497RT, BioRad), dilution 1:200) was applied simultaneously with rabbit anti-mouse M2 marker CD206 (Cat# PA5-101657) or rabbit anti-mouse M1 marker CD86 (Cat# PA5-114995) (both Invitrogen, dilution 1:200). The following day, F4/80 was conjugated using goat anti-rat biotin (Vector Laboratories, with CD206 - dilution 1:200, with CD86-dilution 1:300) followed by Streptavidin 488 (eLIFE, dilution 1:100). CD206 was conjugated using goat anti-rabbit biotin (1:200 Vector Laboratories) followed by Streptavidin 546 (1:1000 Invitrogen). CD86 samples underwent application of secondary antibody donkey anti-rabbit IgG Alexa fluor 594 (1:1000, Invitrogen). Cell nuclei were stained using 4′,6-diamidino-2-phenylindole (DAPI) (1:5000, Sigma-Aldrich). Mouse or Rabbit immunoglobulin G (IgG) was used as the negative control at the same concentration as the primary antibodies.

#### Assessment of Leucocyte Infiltration using QuPath Software

Multiplexed fluorescence images from tissue sections were analyzed using QuPath version 0.2.3, an open-source software platform for digital pathology and whole-slide image analysis (available at https://qupath.github.io). QuPath’s cell segmentation tools were used to detect thousands of individual cells, identify them hierarchically within annotated regions and quantify both cellular morphology and biomarker expression. Immunofluorescent slide imaging was performed using a spinning disk confocal microscope (BC43, Dragonfly 200, Oxford Instruments). To identify macrophage populations, F4/80 was used as a pan-macrophage marker, recognizing a 160 kDa glycoprotein highly expressed by murine macrophages with limited expression in monocytes. CD86 was used to identify M1 pro-inflammatory macrophages, while CD206 (mannose receptor) was used to label M2 homeostatic macrophages. For each animal, the entire uterine cross-section was used as the analysis region. Uterine tissue regions were masked to distinguish decidua and myometrium, and cell counts were performed in each fluorescence channel corresponding to the specific antibody. QuPath reports annotation and detection areas in square micrometres (μm²). For each masked compartment, the number of positive cells was divided by the measured compartment area and scaled to square millimetres, where 1 mm² equals 1,000,000 μm². Macrophage density is therefore reported as the number of positive cells per mm² for each marker.

#### RNA Sequencing and Analysis

Myometrial RNA (collected as described for RT-qPCR, n=3, minimum RINs > 8) from each group (saline, BSCI, IL-1α, and IL-1α/BSCI) at 24 hr was submitted for RNA library preparation (NEB Ultra II Directional polyA mRNA) and sequencing (The Centre for Applied Genomics, Sick Kids, Toronto) on a 10B flowcell. Samples were paired-end, 150 base-pairs in length, and a minimum of 45 million reads were sequenced. Raw reads were quality checked using FastQC (https://www.bioinformatics.babraham.ac.uk/projects/fastqc/), adaptors and poor quality reads removed, and trimming completed using TrimGalore! (https://github.com/FelixKrueger/TrimGalore). Trimmed reads were then aligned to GRCm38 using Hisat2 (67). For visualization, Bedgraphs were generated from bam files using bamCoverage with TPM normalisation and uploaded to the UCSC genome browser (genome.ucsc.edu). Reads were quantified using Salmon, controlling for GC bias (68), and differential reads were identified using DESeq2 (69). Volcano plot visualizations for differentially expressed genes (abs. fold change ≥ |2|, FDR < 0.05) were generated using ggplot2. Gene Ontology (GO) enrichment was conducted on genes significantly increasing or decreasing expression using g:Profiler (70). Data are available at GEO repository (GSE317954).

#### Assay for Transposase-Accessible Chromatin with Sequencing (ATAC-seq) of Myometrial Tissue

Myometrial tissues (n=3/condition) collected and crushed in liquid nitrogen from saline, BSCI, IL-1α, and IL-1α/BSCI groups at 24 hr and submitted to the Princess Margaret Genomics Centre (PMGC), Toronto, Canada, for ATAC-seq. Sample libraries were prepared following the Omni ATAC-seq protocol (71). Samples were sequenced on the Illumina NovaSeq X, which were 50 bp in length and paired-end. Sequencing data files were submitted to and are publicly accessible at the GEO repository (GSE292301).

#### Read Alignment and Peak Calling for ATAC-seq

Prior to processing, raw reads were analyzed for quality using FastQC (https://www.bioinformatics.babraham.ac.uk/projects/fastqc/). Raw reads were then analyzed using the ENCODE ATAC-seq pipeline (https://github.com/ENCODE-DCC/atac-seq-pipeline). Trimmed reads from the ENCODE ATAC-seq pipeline were then re-checked for quality using FastQC to verify trimming parameters and maintain/improve quality post-trimming before proceeding. As part of the ENCODE ATAC-seq pipeline, reads were trimmed using cutadapt (72) and aligned to the mm10 genome with aligner Bowtie2 (73). Peaks were called using Model-based Analysis of ChIP-Seq (MACS)2 (narrowpeaks setting;(74)) and significantly conserved peaks between replicates identified using Irreproducible Discovery Rate (IDR;(75)). To identify differentially accessible chromatin regions between conditions, .bam and .narrowpeak files from each replicate (n=3) were read into DiffBind R Package, and blacklists applied (76). Using all reads, correlation coefficients (plot) and principal component analyses (dba.plotPCA) were conducted. Contrasts were established, and DESeq2 was used to identify all regions of differential chromatin accessibility (FDR ≤ 0.05) in each contrast. To generate BedGraphs for data visualization bam files generated using the ENCODE ATAC-seq pipeline for each biological replicate were merged and sorted using Samtools. Merged files were processed using bamCoverage from Deeptools) (77) and uploaded to the UCSC genome browser (genome.ucsc.edu) for visualization. Similarly, .narrowpeak files (from MACS2 applied by the ENCODE ATAC-seq pipeline) containing regions identified as peaks were also uploaded, appearing as bars under .bedGraph tracks. Volcano plot visualizations for differentially accessible chromatin (abs. fold change ≥ |2|, FDR < 0.05) was generated using ggplot2.

#### Statistical Analysis

The normality of datasets was determined by the Shapiro–Wilk test. The Grubbs’s outliers test was implemented with the assumption of a normally distributed population. Mann–Whitney test (two-tailed), Pearson correlation, Chi-square analysis, Fisher’s exact test, Kaplan–Meier analysis (Log-rank), paired two-tailed Student’s t-test, or unpaired two-tailed Student’s t-test (two groups) or two-way ANOVA (more than two groups) were used as appropriate. The data are presented as the means ± SEMs. The significance level was set at P < 0.05 (*), P < 0.01 (**), and P < 0.001 (***).

## RESULTS

### BSCI Reduces the Incidence of IL-1α-induced PTB

Using the pregnant mouse as a model, we investigated whether BSCI could prevent or delay PTB induced by ultrasound-guided intra-amniotic injections of IL-1α (Fig. 2A). No delivery occurred for the first 12 hrs after administration of IL-1α (400 ng/amniotic sac); however, all treated dams (n = 8) delivered within 24 hrs, with a mean ‘injection-to-delivery interval’ of 23.7 ± 4.4 hrs (**Fig. 2B, C**). No maternal mortality or morbidity was observed. Control mice receiving daily intra-amniotic saline injections delivered at term (GD18.5-20.5, n=6). Similarly, dams receiving daily injections of FX125L alone (‘BSCI-only’ group, n=9) delivered at term, with no apparent adverse outcomes for the dam or pups (Fig. 2B, C). Pretreatment with BSCI (10 mg/kg/day) significantly reduced the incidence of PTB following IL-1α challenge, from 100% to 35.7% (5 of 14 dams delivered preterm, P < 0.01, **Fig. 2C**). BSCI extended the mean treatment-to-delivery interval from 23.7 ± 4.4 to 50.6 ± 26.2 hrs (**Table 1 and Fig. 2B**). The mean number of pups delivered per litter did not differ significantly between the vehicle (saline), IL-1α, BSCI, and IL-1α/BSCI groups (**Table 1**).

**Figure 2.**
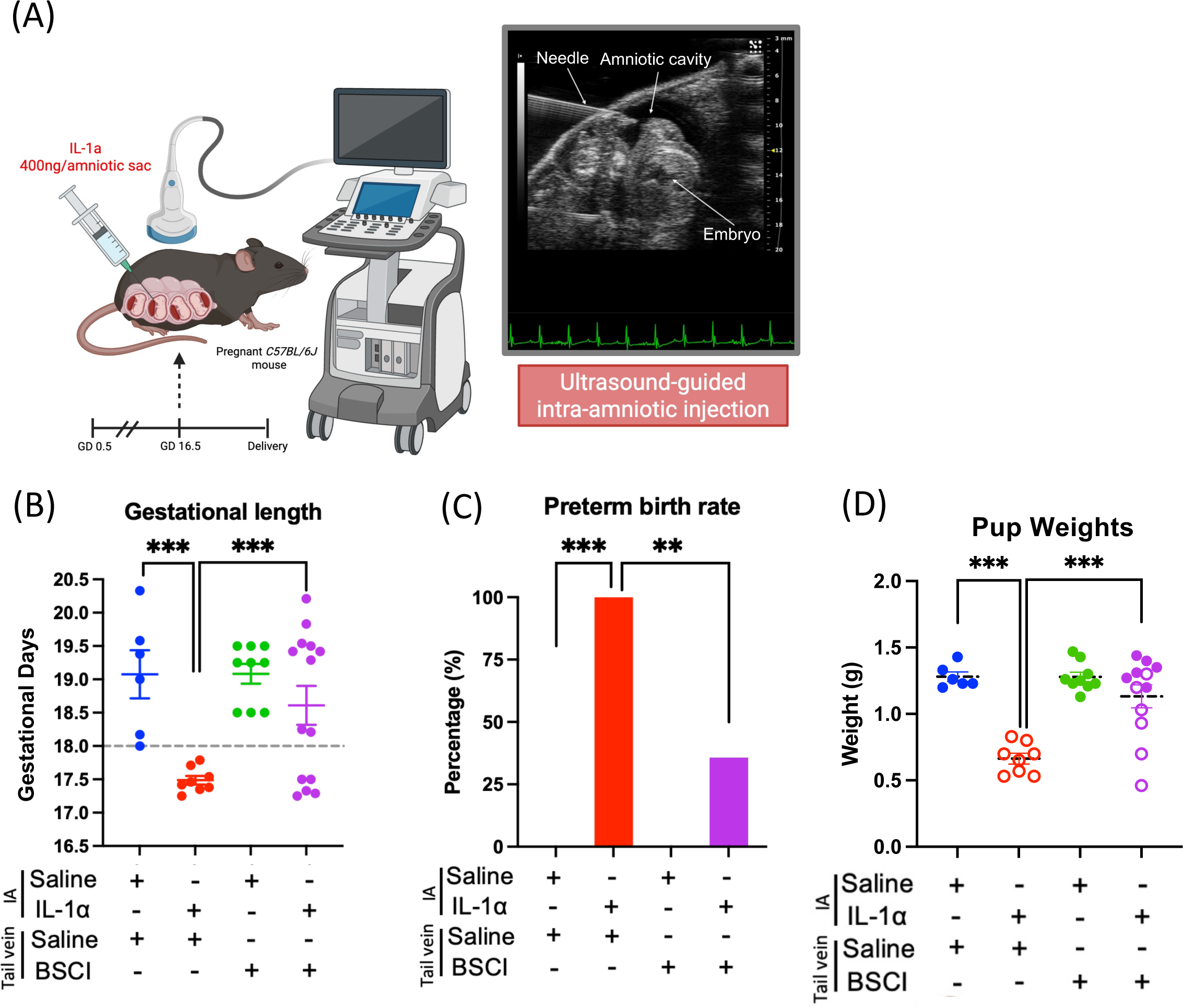
The BSCI reduces the incidence of preterm birth and neonatal mortality induced by in utero sterile inflammation. **(A)** To induce *in utero* sterile inflammation, the alarmin IL-1α (400 ng/amniotic sac) was intra-amniotically administered to C57BL/6 dams under ultrasound guidance on gestational day (GD) 16.5. Dams were monitored until delivery, and neonatal survival and weight were recorded. Treatments are indicated below each panel (IA, intra-amniotic injection; tail vein, daily intravenous injection): saline/saline (n = 6), IL-1α/saline (n = 8), saline/BSCI (n = 9), IL-1α/BSCI (n = 14). **(B)** Gestational length is shown for each treatment group in gestational days (GD). The dashed line indicates the threshold of term delivery after GD 18. **(C)** Preterm birth rates are shown as bar plots. **(D)** Fetal weights at delivery are shown as litter means. For term litters, each value is the average of 3 pups per dam; in the 2 term litters with intrauterine fetal demise, only normally developed pups were weighed, so these values exclude the undeveloped pups. For preterm litters, in which partial cannibalization precluded standardized sampling, each value is the mean of the recoverable pups. Open symbols denote litters without live-born pups. The IL-1α/BSCI group shows 12 of 14 litters: one litter was fully cannibalized, so no pup weights could be recorded, and one litter mean was excluded as a statistical outlier (Grubbs test). Gestational length and pup weights were analyzed by two-way ANOVA followed by a Bonferroni post-test; preterm birth rates were compared by Fisher’s exact test. Results were expressed as mean ± SEM. Significant differences are indicated by ** (P < 0.01) and *** (P < 0.001).

**Table 1.** BSCI reduces the incidence of IL-1α-induced PTB and allows animals to deliver at term.

| Treatment | N | Preterm<br>birth rate<br>(%) | Term<br>birth rate<br>(%) | Treatment<br>to delivery<br>(h) | Mean N<br>of pups/<br>litter | Neonate/fetal<br>weight (g) | Placental<br>weight (g) |
| --- | --- | --- | --- | --- | --- | --- | --- |
| Vehicle | 6 | 0/6 (0) | 6/6 (100) | 67.8±17.5 | 7.8 | 1.28±0.08 | -- |
| IL-1α | 8 | 8/8 (100) | 0/8 (0) | 23.7±4.4 | 7.7 | 0.66±0.12<sup>#</sup> | 0.09±0.04<sup>§</sup> |
| BSCI | 9 | 0/9 (0) | 9/9 (100) | 68±3 | 8.1 | 1.28±0.11 | 0.15±0.06 |
| IL-1α+BSCI | 14 | 5/14<br>(35.7) | 9/14<br>(64.3) | 50.6±26.2 | 7.2 | 1.13±0.30 | 0.18±0.22 |

<sup>#</sup>Indicates the difference between the IL-1α group and the vehicle and IL-1α +BSCI groups (P < 0.001).
<sup>§</sup>The delivery time in IL-1α group is GD17, while in BSCI or IL-1α +BSCI groups – GD19-20.
Neonate/fetal weights are litter means (average of 3 pups per dam). In term litters with fetal demise, litter means include normally developed pups only. The IL-1α +BSCI value covers 12 of 14 litters; one litter was cannibalized by the dam and one litter mean (2.2 g) was excluded as a statistical outlier (Grubbs test).

**Table 2.**
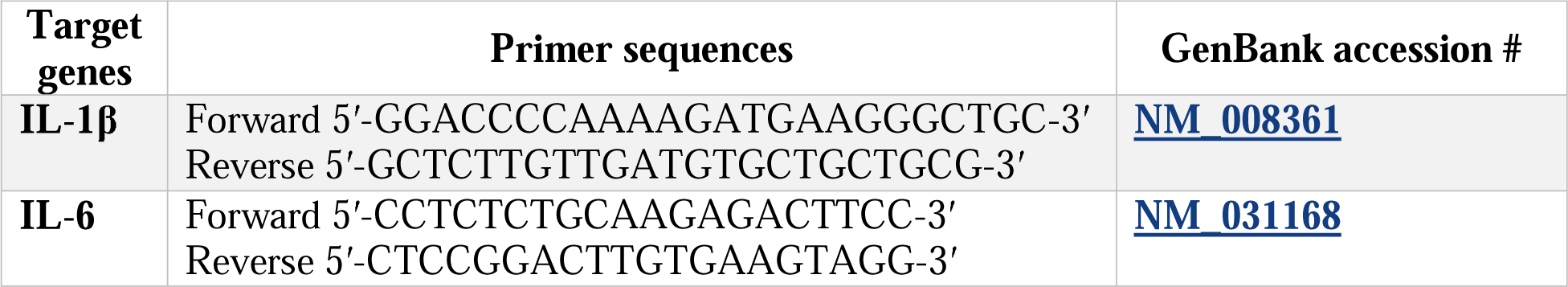
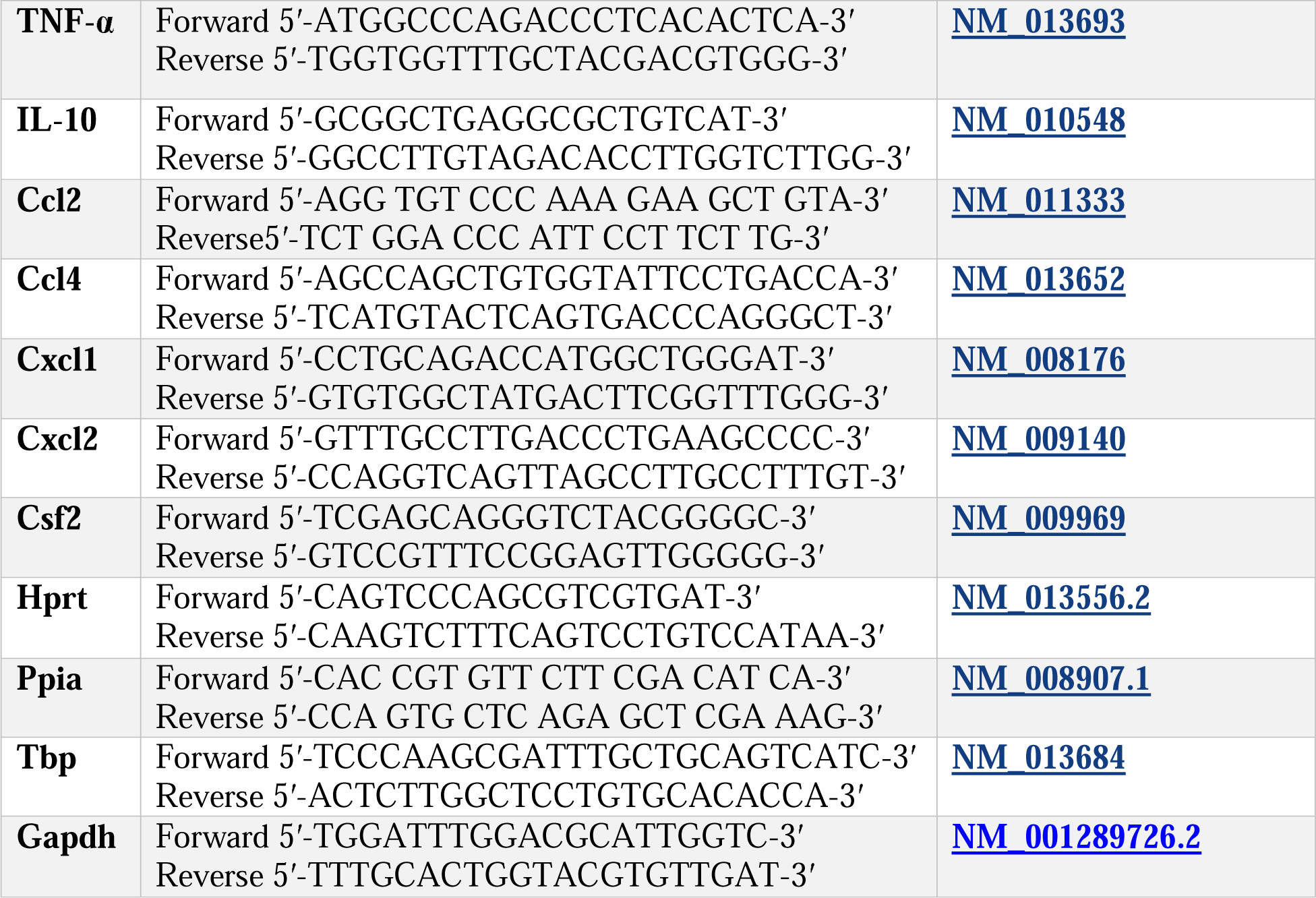
Real-time PCR primer sequences of a panel of genes involved in inflammatory response and housekeeping genes.

We next assessed fetal and placental weights to further characterize the outcomes of BSCI treatment. Fetal weights were compared among the BSCI-only control group, which received daily BSCI injections from GD15.5 until term (n = 9); the vehicle group, which received saline (n = 6); and the IL-1α/BSCI group (n = 14) (Table 1; Fig. 2D). Among the 14 dams pretreated with BSCI before IL-1α challenge, 5 delivered preterm. None of the pups in these litters survived, and the dams cannibalized the pups, precluding individual pup weight measurements. The corresponding litter weights were below those of litters with live-born pups, consistent with delivery at an earlier gestational age (Fig. 2B, D). The remaining 9 dams delivered at term. Live pups were observed in 7 of these 9 litters, whereas the other 2 litters contained no pups showing vital signs at delivery; most pups appeared undeveloped, consistent with intrauterine fetal demise; in these 2 litters, only the normally developed pups were weighed, so their litter means in Fig. 2D exclude the undeveloped pups. Placental weights in the IL-1α/BSCI group were highly variable (0.18 ± 0.22 g; **Table 1**), with the group mean inflated by a single abnormal 0.68 g placenta from a term litter without vital signs. Thus, although BSCI pretreatment significantly reduced the incidence of PTB and prolonged pregnancy following IL-1α challenge, fetal rescue remained incomplete.

### BSCI Prevents IL-1α-induced Cytokine Expression in Maternal and Fetal Tissues

To investigate whether IL-1α-induced IAI could activate the immune system of pregnant mice, we measured mRNA and protein expression of numerous pro- and anti-inflammatory cytokines in maternal plasma and liver. Within 2 hrs of administration, IL-1α significantly (P < 0.001) increased plasma levels of IL-1α, IL-6, IL-10, IL-12p70, TNF-α, and CSF2 as compared to the saline-injected animals (**Fig. 3**, n = 6/group), indicating systemic maternal inflammation. All cytokine proteins fell to control levels by the 24 hrs time-point. Importantly, BSCI significantly (P < 0.001) attenuated the increase in IL-1α, IL-6, IL-10, IL-12p70, and CSF2 protein plasma levels at 2 hrs after IL-1α injection; TNF-α level was also decreased, however, non-significantly (P = 0.192, **Fig. 3**). BSCI pretreatment alone did not affect plasma cytokine levels compared to the vehicle control group.

**Figure 3.**
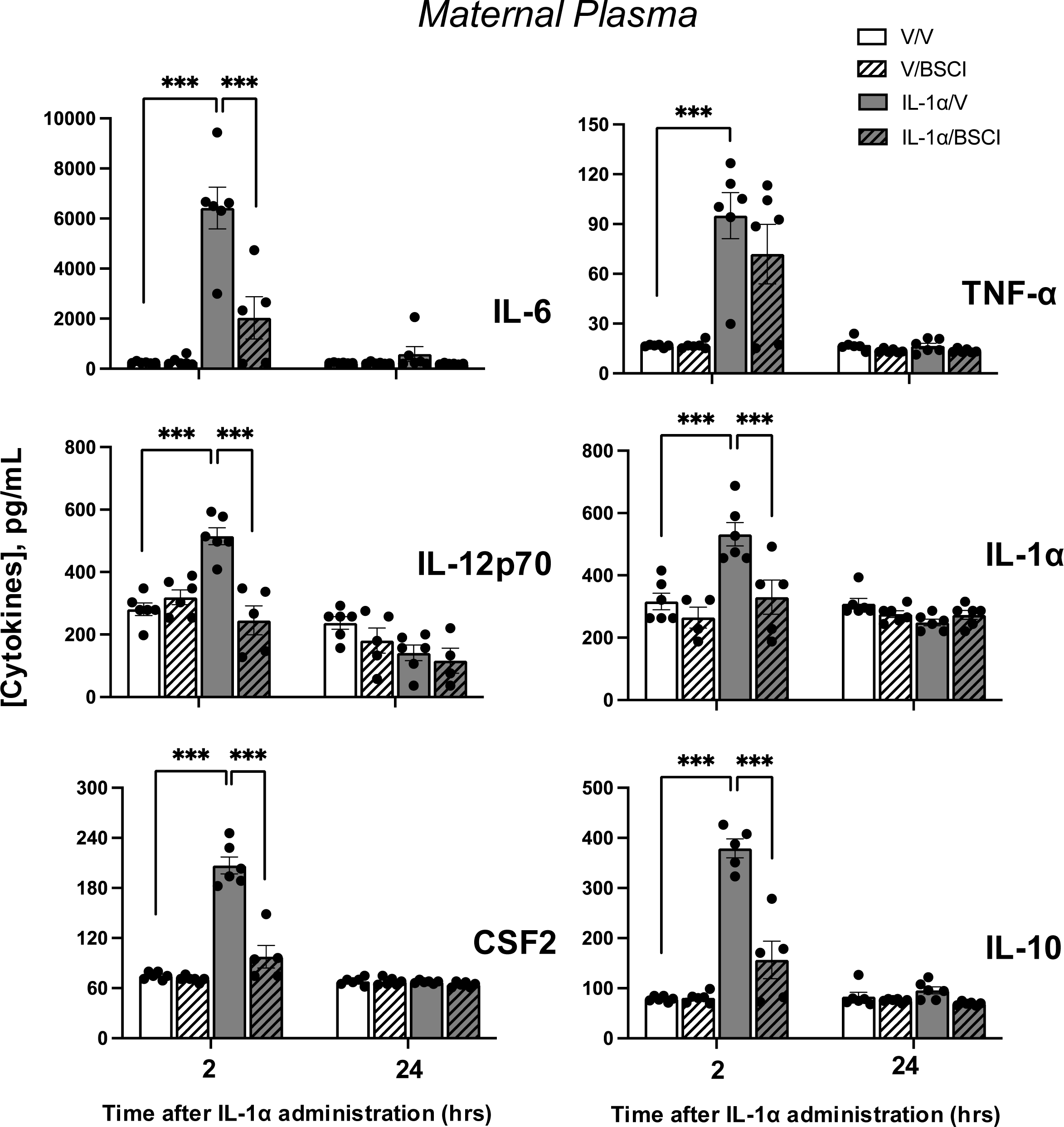
Temporal changes in cytokine protein levels of the maternal blood plasma from GD16.5 and 17.5 pregnant mice challenged with IL-1α and treated with Broad Spectrum Chemokine Inhibitor (BSCI). Pro-inflammatory (IL-1α, IL-6, IL-12p70, TNF-α, CSF2) and anti-inflammatory (IL-10) cytokines; protein expression was detected by multiplex magnetic bead assay following local IL-1α administration and pretreatment with BSCI. Shown are vehicle (white bars), BSCI-treated (striped bars), IL-1α-injected (grey bars), and IL-1α-injected BSCI-treated samples (striped grey bars), n =5–6/group. Two-way ANOVA was utilized, followed by a Bonferroni post-test. Results were expressed as mean ± SEM. Significant difference between groups is indicated by *** (P < 0.001).

We found evidence of a systemic maternal inflammatory response after intraamniotic IL-1α injection. Expression of genes encoding pro-inflammatory (*Il6, IL1*β*, Tnf*α), anti-inflammatory (*Il10*) cytokines, and chemokines *Ccl2/*Mcp1 (major chemoattractant for monocytes), *Cxcl1/*KC/Groα (major chemoattractant for neutrophils), and *Cxcl2*/Mip2a were increased in maternal liver (P < 0.01-0.001) 2 hrs after IL-1α injection (5-to 180-fold increase, **Fig. 4A,B**). This increase in maternal liver inflammatory gene expressions was largely prevented by BSCI (**Fig. 4A,B**).

**Figure 4.**
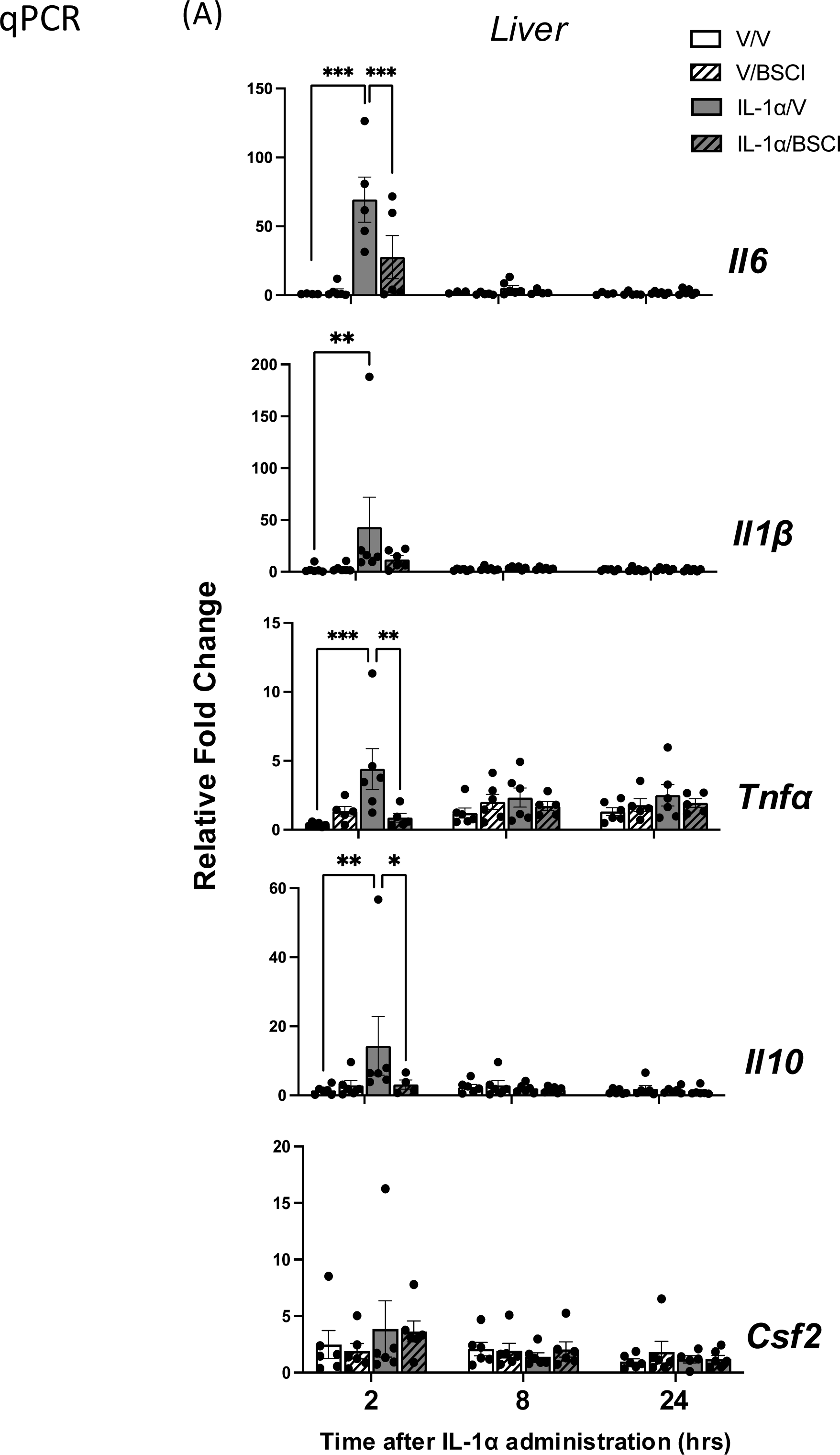
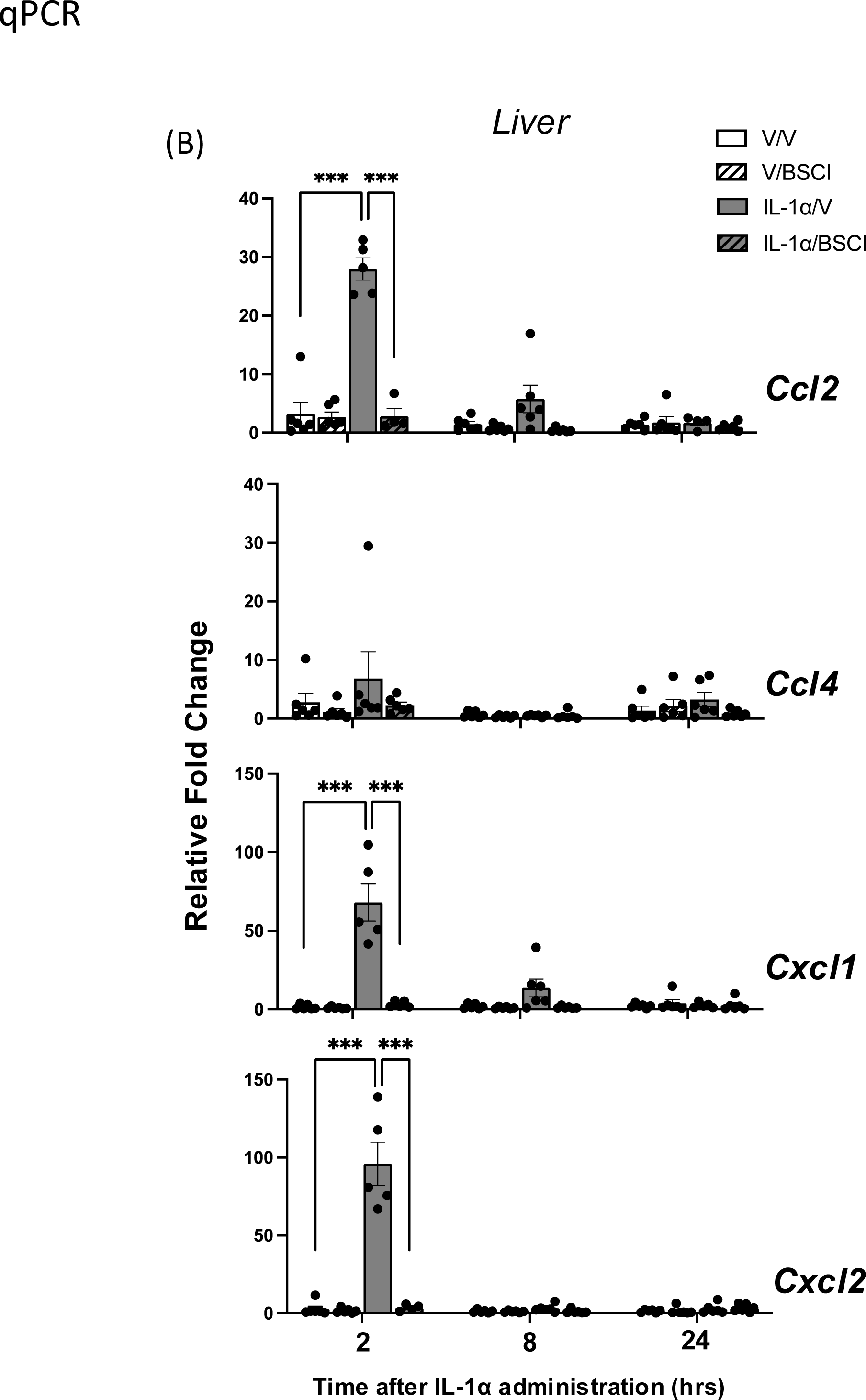
Changes in cytokine mRNA in the mouse liver of GD16 and 17 pregnant mice following local IL-1α intra-amniotic infusion and pretreatment with Broad Spectrum Chemokine Inhibitor (BSCI). **(A)** Pro-inflammatory (*Il6*, *Il1*β, *Tnf*α, *Csf2/Gm-scf*) and anti-inflammatory (*Il10*) cytokines; **(B)** Chemokines *Ccl2/Mcp1*, *Ccl4/Mip1*β, *Cxcl1/KC/Gro*α, and *Cxcl2/Mip2*α transcripts were detected by Real-Time RT-qPCR in mouse liver. Shown are samples from vehicle (white bars), BSCI-treated (striped bars), IL-1α-injected (grey bars), and IL-1α-injected + BSCI-treated pregnant animals (striped grey bars), n = 5–6/group. Two-way ANOVA was utilized, followed by a Bonferroni post-test. Results were expressed as mean ± SEM. Significant difference is indicated by * (P < 0.05), ** (P < 0.01) and *** (P < 0.001).

Next, we examined the direct effect of BSCI on IL-1α-induced IAI by studying mRNA expression and protein production of pro- and anti-inflammatory cytokines and chemokines in uterine (myometrium and decidua) and fetal (placenta and amniotic fluid) tissues. All inflammatory genes studied by RT-qPCR were significantly increased in dams at 2 hrs post IL-1α compared to saline-injected animals (myometrium: 2-to 1000-fold increase, **Fig. 5**, and decidua: 2-to 760-fold increase, **Fig. 6**; P < 0.001 for all), but fell to baseline levels by 8 hrs after IL-1α administration. BSCI pretreatment significantly (P < 0.05-0.001) attenuated IL-1α-induced expression of cytokine or chemokine mRNA at 2 hr time-points (*Il6, Tnf*α*, Il1*β*, Il10*, *Csf2* (**Fig. 5A**); chemokines *Ccl2, Ccl4, Cxcl1, Cxcl2* (**Fig. 5B**); as well as of transcripts for pro-labor contraction-associated proteins *Nfkb1, Ptgs2, Akr1c18, Gja1* (**Fig. 5C**).

**Figure 5.**
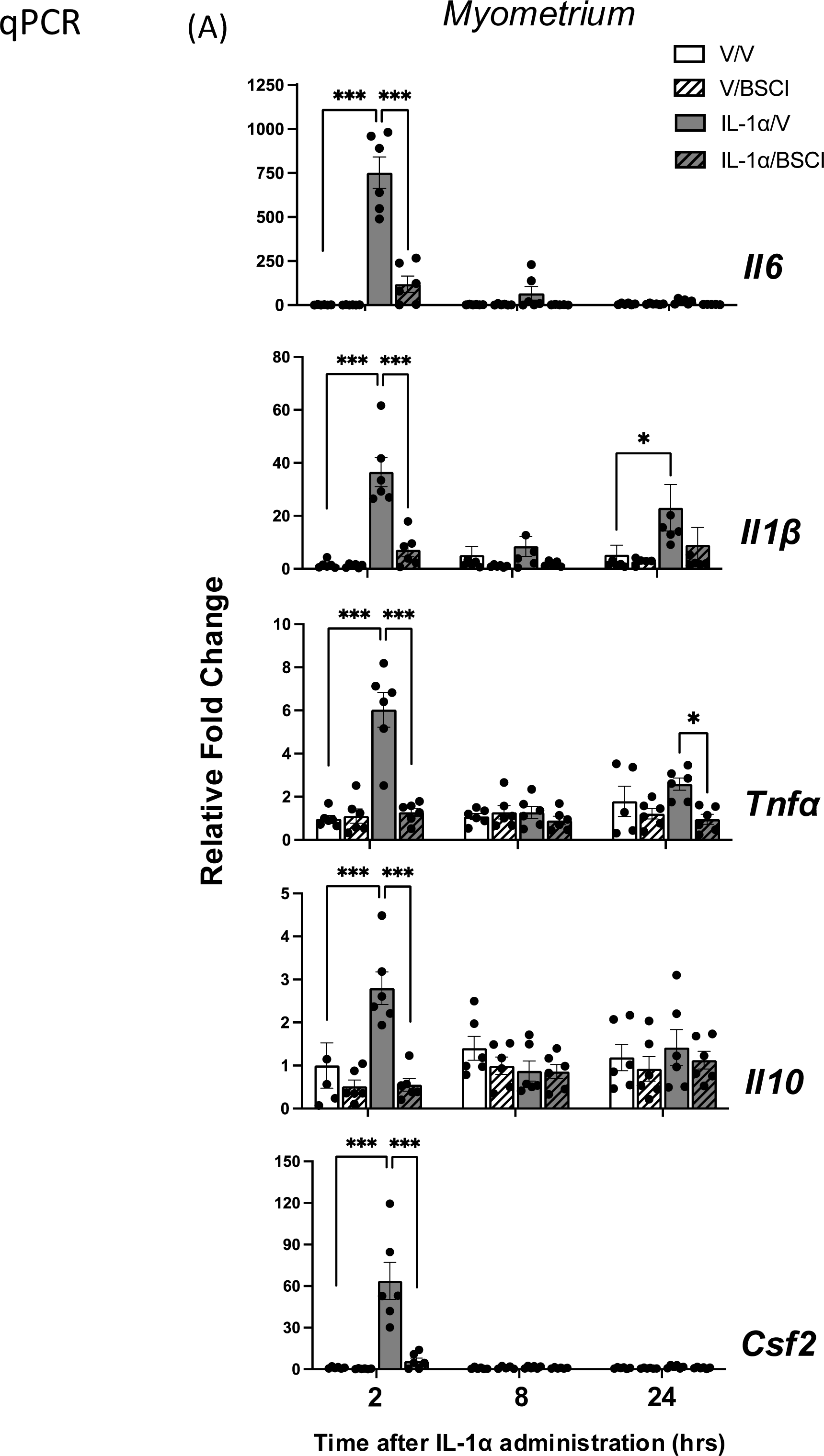
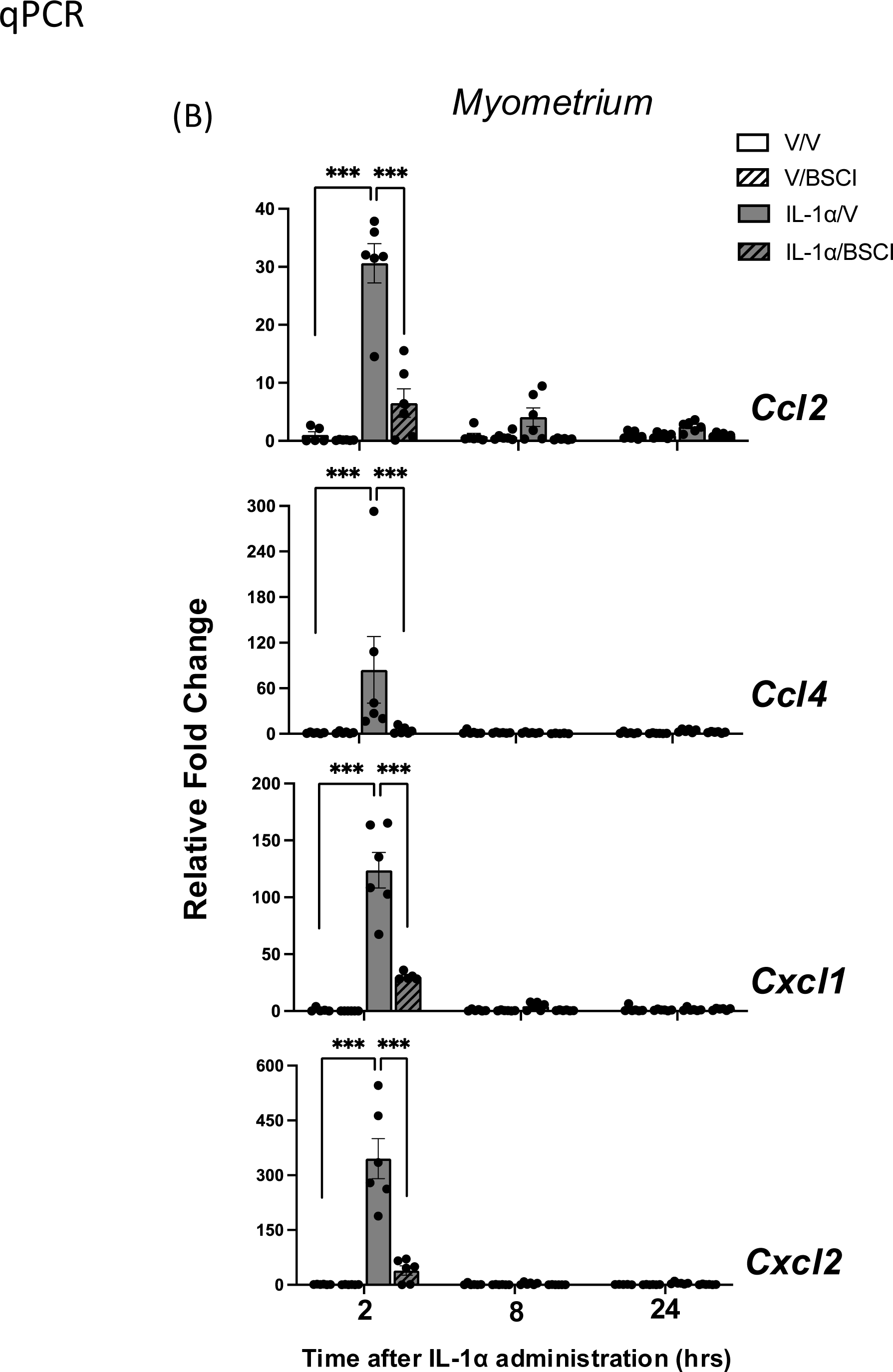
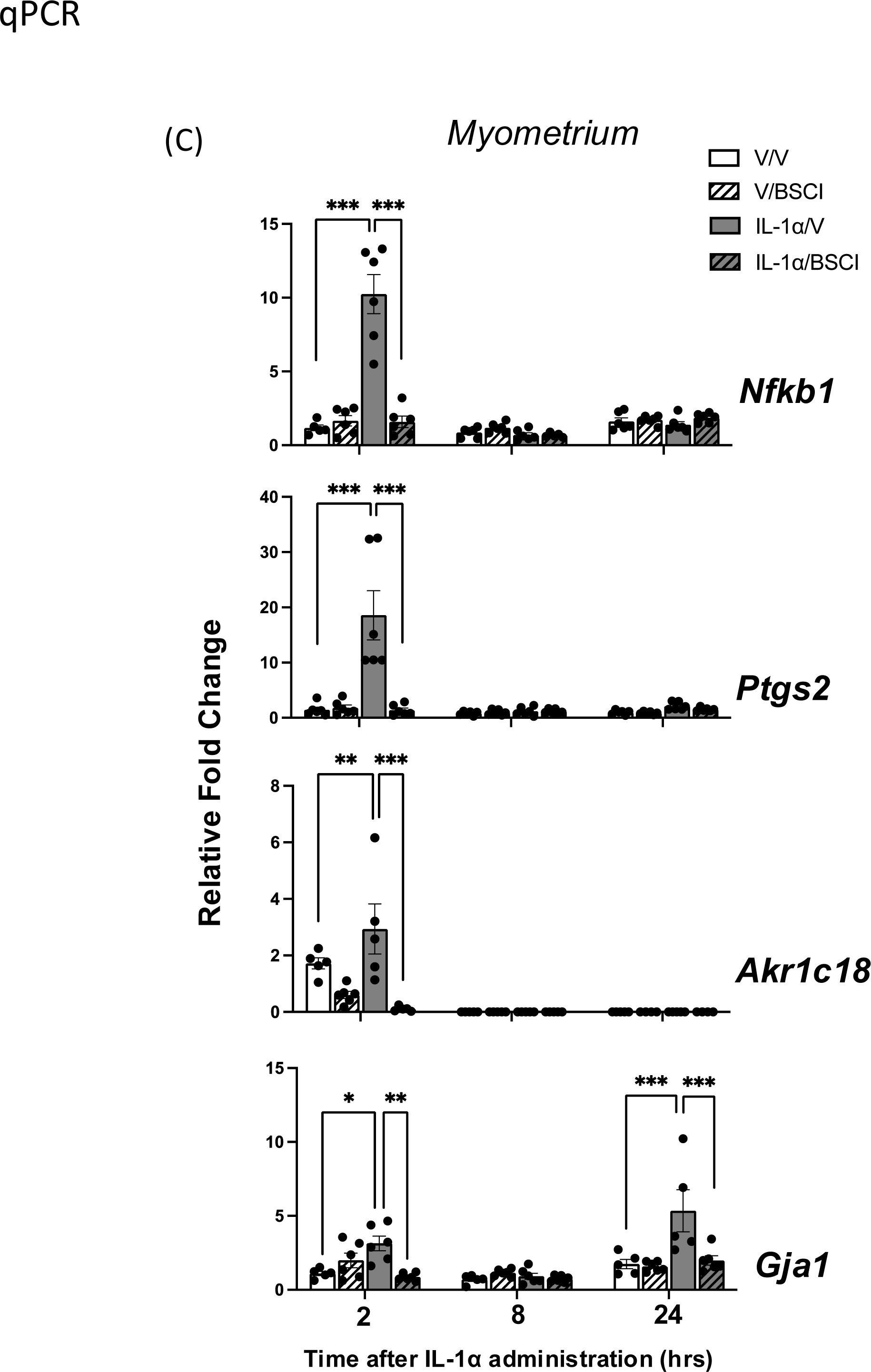
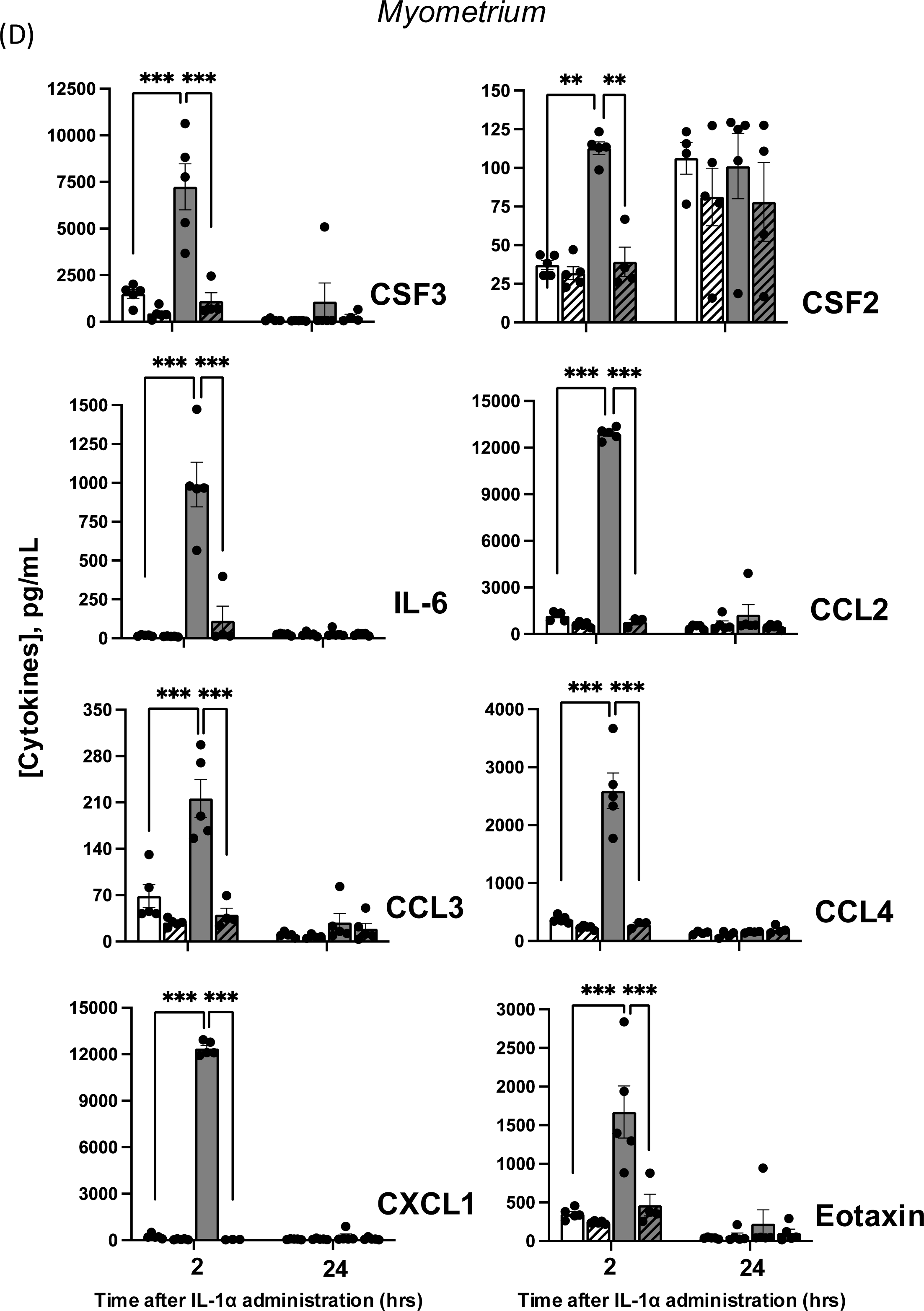
Changes in cytokine mRNA and protein levels in the mouse myometrium of GD16 and GD17 pregnant mice following local IL-1α intra-amniotic infusion and pretreatment with Broad Spectrum Chemokine Inhibitor (BSCI). **(A)** Transcript levels of pro-inflammatory (*Il6*, *Il1*β, *Tnf*α, *Csf2/Gmscf*) and anti-inflammatory (Il10) cytokines;**(B)** Chemokines (*Ccl2/Mcp1*, *Ccl4/Mip1*β, *Cxcl1/KC/Gro*α and *Cxcl2/Mip2*α); and **(C)** Pro-labor genes *Nfkb1*, *Ptgs2/Cox2*, *Akr1c18/20*α*HSD*, and *Gja1/Cx43* detected by Real-Time RT-PCR. **(D)** Protein levels of pro-inflammatory cytokines (CSF2, CSF3, IL-6) and chemokines (CCL2, CCL3, CCL4, CXCL1, Eotaxin) were detected by multiplex magnetic bead assay (BioRad). Shown are vehicle (white bars), BSCI-treated (striped bars), IL-1α-injected (grey bars), and IL-1α-injected BSCI-treated samples (striped grey bars), n = 5–6/group. Two-way ANOVA was utilized, followed by a Bonferroni post-test. Results were expressed as mean ± SEM. Significant difference is indicated by * (P < 0.05), ** (P < 0.01), and *** (P < 0.001).

**Figure 6.**
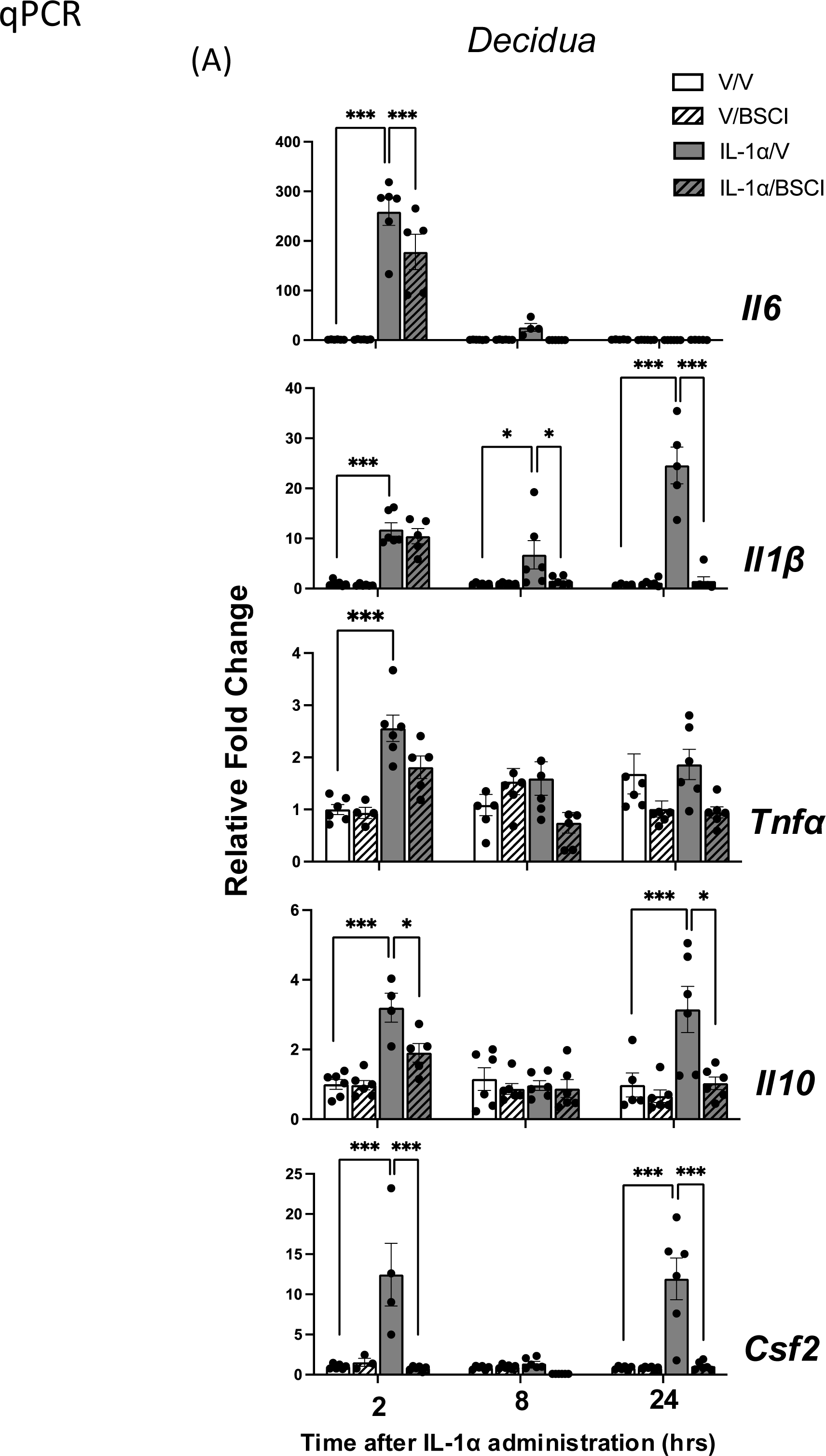
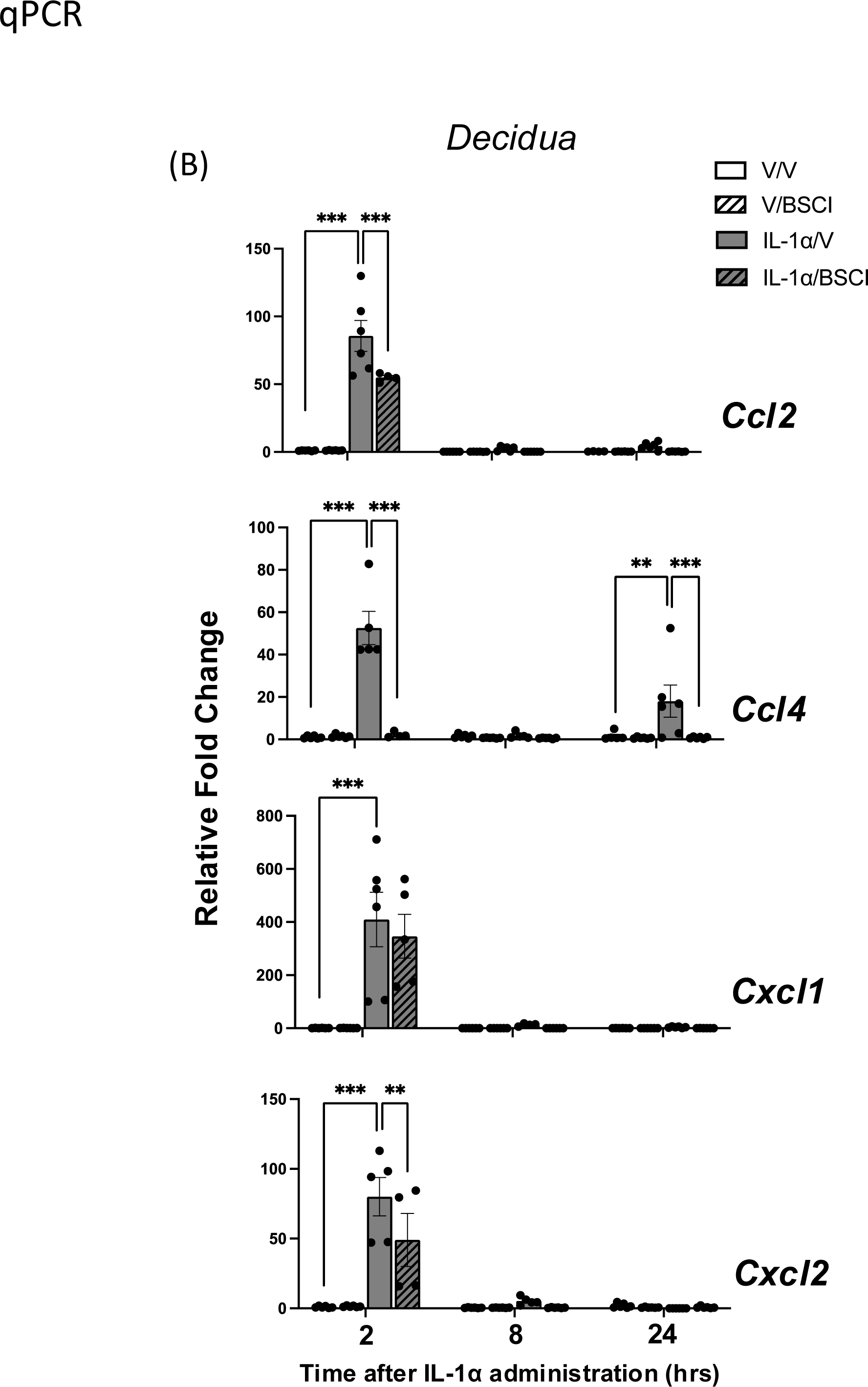
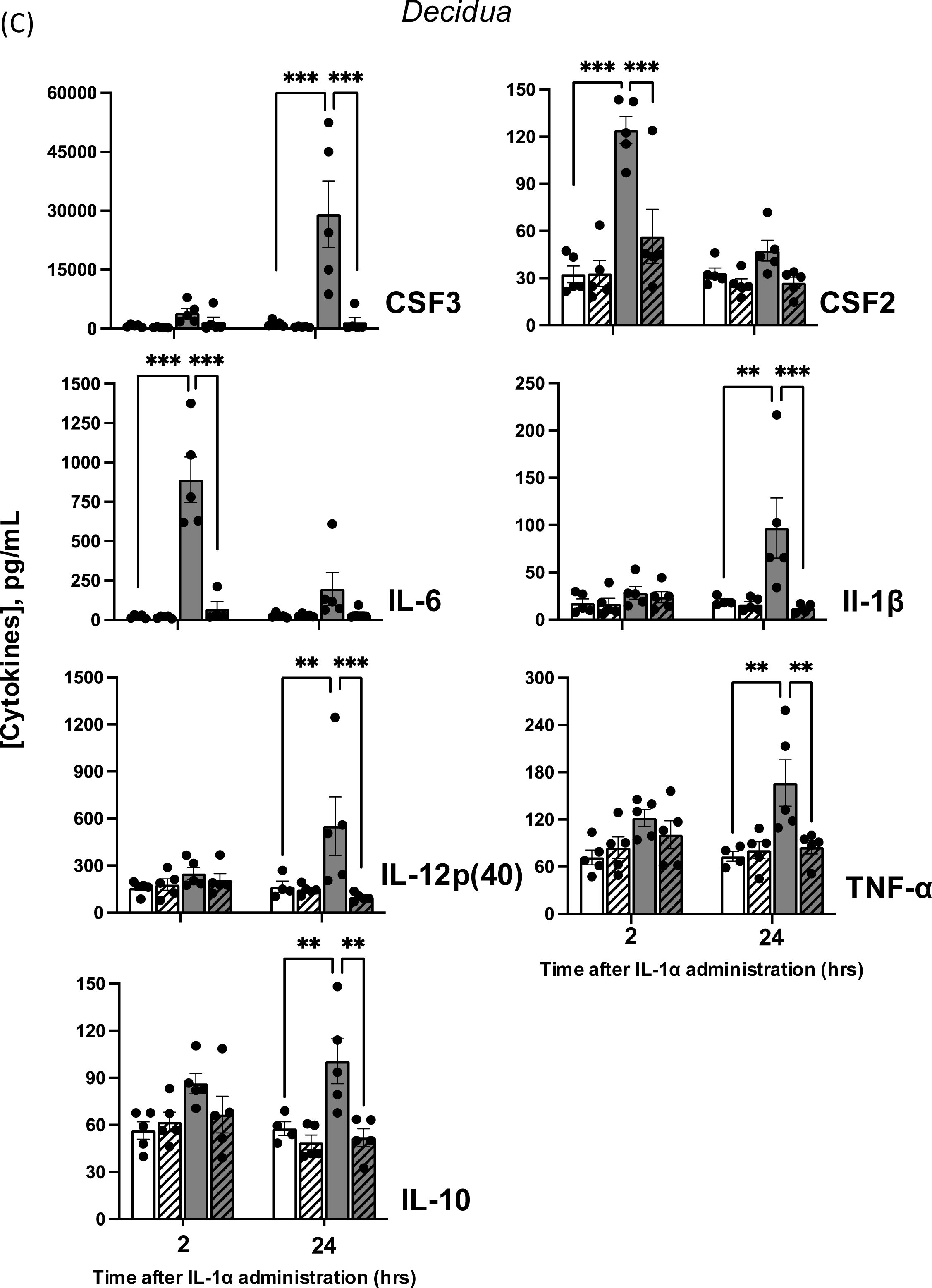
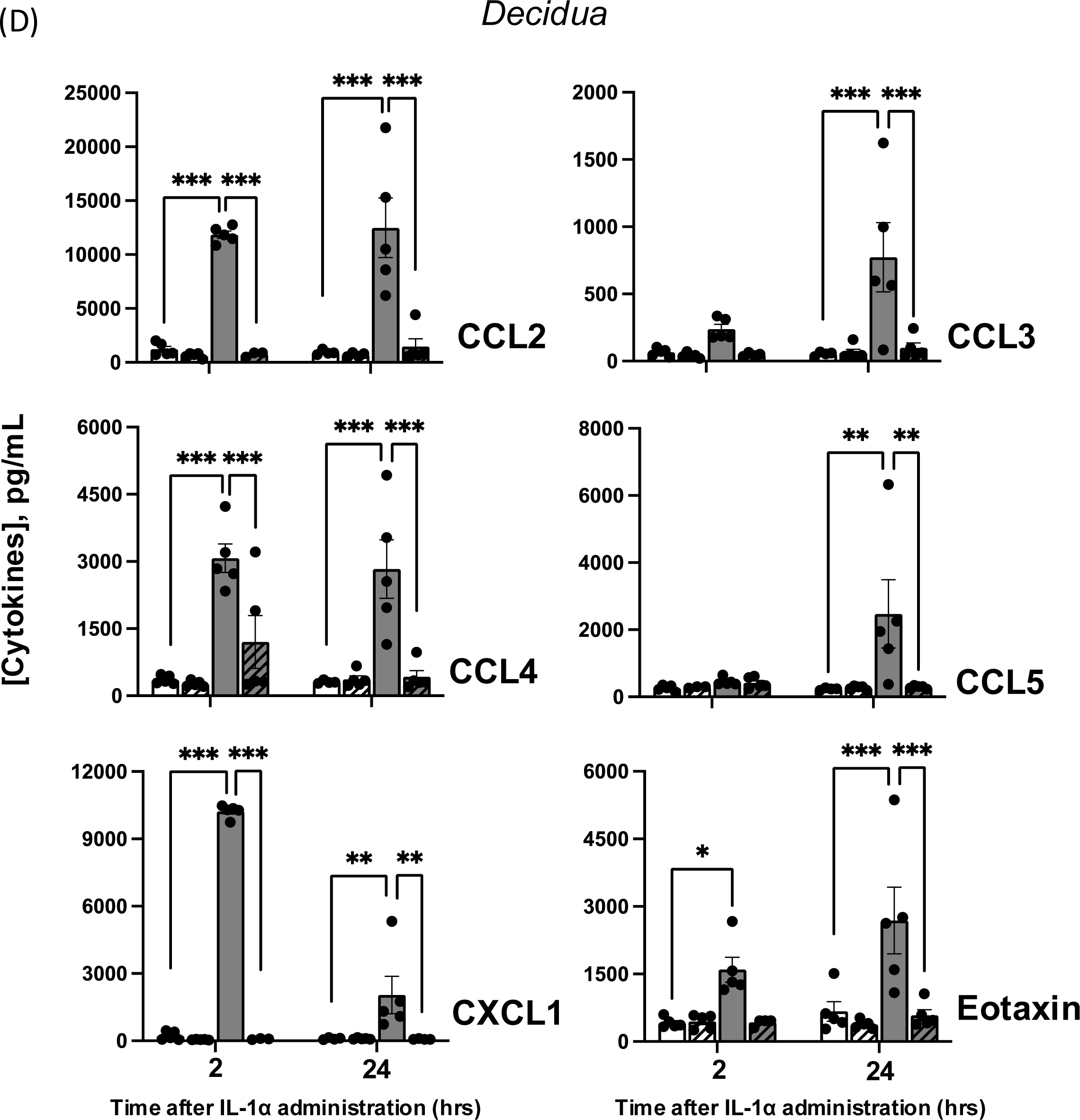
Changes in cytokine mRNA and protein levels in the decidua of GD16 and GD17 pregnant mice following local IL-1α intra-amniotic infusion and pretreatment with Broad Spectrum Chemokine Inhibitor (BSCI). **(A)** Transcript levels of pro-inflammatory (*Il6*, *Il1*β, *Tnf*α, *Csf2*) and anti-inflammatory (*Il10*) cytokines; **(B)** Chemokines (*Ccl2/Mcp1*, *Ccl4/Mip1*β, *Cxcl1/KC/Gro*α and *Cxcl2/Mip2*α) were detected by Real-Time RT-qPCR. **(C)** Protein levels of pro-inflammatory (CSF2, CSF3, IL-6, IL-12(p40), IL-1β, TNF-α), anti-inflammatory (IL-10) cytokines; and **(D)** chemokines (CCL2, CCL3, CCL4, CCL5, CXCL1, Eotaxin) were detected by multiplex magnetic bead assay. Shown are vehicle (white bars), BSCI-treated (striped bars), IL-1α-injected (grey bars), and IL-1α-injected BSCI-treated samples (striped grey bars), n=5–6/group. Two-way ANOVA was utilized, followed by a Bonferroni post-test. Results were expressed as mean ± SEM. Significant difference is indicated by * (P < 0.05), ** (P < 0.01), and *** (P < 0.001).

We found similar changes in the decidua, with intra-amniotic IL-1α inducing a rapid 2 hr increase in inflammatory gene expression, which was attenuated by pretreatment with BSCI (**Fig. 6A**, **6B**). Importantly, 24 hrs after IL-1α injections, at the time of preterm labor, transcript levels of three cytokines, the major pro-inflammatory cytokine *Il1*β, anti-inflammatory cytokine *Il10*, and *Csf2,* which supports leukocyte production, survival, and activation within the decidua, all remained elevated (P < 0.001), suggesting an active role of decidual inflammation during PTB (**Fig. 6A**).

As with gene expression, intra-amniotic IL-1α injection produced a transient 2 hr, significant (P < 0.01-0.001) increase in myometrial cytokines IL-6, CSF2/GM-CSF, CSF3/G-CSF, and chemokines CCL2, CCL3, CCL4, CXCL1, and Eotaxin as compared to vehicle (saline), an effect that was attenuated by BSCI pretreatment (P < 0.01-0.001) (**Fig. 5D**). In the uterine decidua IL-1α induced a significant (P < 0.001) increase in cytokines IL-6, CSF2 and chemokines CCL2, CCL4, CXCL1, and Eotaxin at 2 hrs which was sustained through to preterm labor for the cytokines CSF3, IL-1β, IL-12p(40), TNF-α, IL-10, and chemokines CCL2, CCL3, CCL4, CCL5, and Eotaxin (P < 0.01-0.001) (**Fig. 6C, D**). These data suggest that the decidua is a major driver of inflammation throughout active labor, while in the myometrium, the transient increase in tissue inflammation induced by IL-1α leads to increased expression of contraction-associated genes such as *Gja1*. Nevertheless, pretreatment with BSCI attenuated both the acute and longer-term increases in decidual cytokines/chemokines induced by IL-1α. Thus, these results indicate that the increased levels of uterine cytokines/chemokines driving IL-1α-induced PTB are inhibited by administration of the BSCI compound FX125L.

We next determined the impact of intra-amniotic IL-1α on fetal inflammation. Surprisingly, intra-amniotic IL-1α injection induced a transient increase only in amniotic fluid IL-6 protein levels with no change in TNF-α, CSF2, and IL-10 proteins (IL-1α, IL-1β, IL-2, IL-4, IL-12p70 were not detectable). Again, BSCI pretreatment attenuated IL-1α-induced increase in amniotic fluid IL-6 protein levels (**Fig. 7**). In the fetal placenta, mRNA levels of *Il6, Il1*β*, Tnf*α*, Csf2* cytokines (**Fig. 8A**) and chemokines *Ccl2, Ccl4, Cxcl1, Cxcl2* (**Fig. 8B**) were significantly (P < 0.05-0.001) up-regulated by IL-1α-induced IAI as compared to vehicle (saline). Importantly, expression of all inflammatory genes in the placenta except *Ccl4* was significantly attenuated by the BSCI (IL-1α versus IL-1α + BSCI groups, P < 0.01-0-001, **Fig. 8A-B**). The major anti-inflammatory cytokine *Il10* in the placenta was not affected by either IL-1α or by BSCI after 2 hrs; however, by 24 hrs post-injection, *Il10* and *Il1*β gene expression were increased in all groups, regardless of treatment, as compared to the 2 hrs time point (**Fig. 8A**).

**Figure 7.**
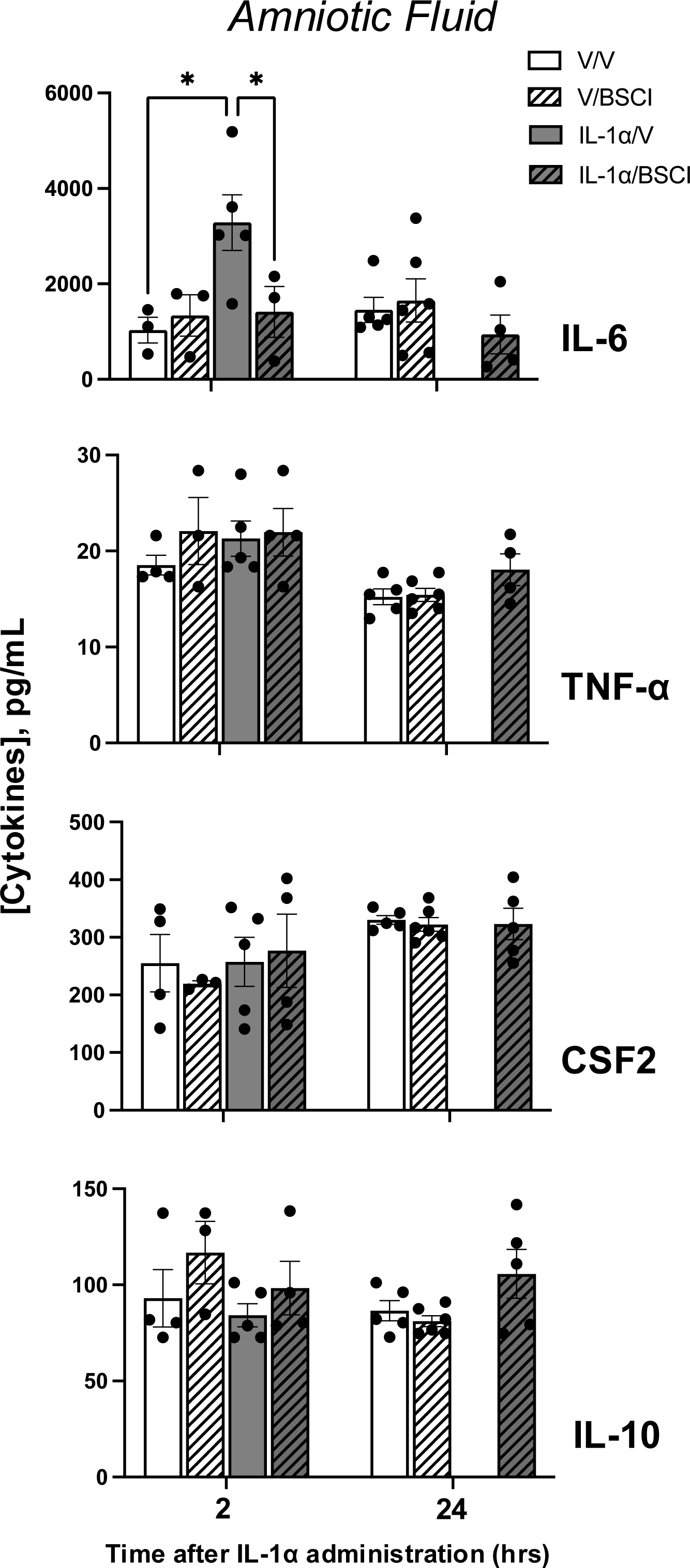
Temporal changes in cytokine protein levels of the amniotic fluid from GD16.5 and 17.5 pregnant mice challenged with IL-1α and treated with Broad Spectrum Chemokine Inhibitor (BSCI). Pro-inflammatory (IL-6, TNF-α, CSF2) and anti-inflammatory (IL-10) cytokines; protein expression was detected by multiplex magnetic bead assay following local IL-1α administration and pretreatment with BSCI. Shown are vehicle (white bars), BSCI-treated (striped bars), IL-1α-injected (grey bars), and IL-1α-injected BSCI-treated samples (striped grey bars), n = 5–6/group. Two-way ANOVA was utilized, followed by a Bonferroni post-test. Results were expressed as mean ± SEM. Significant difference is indicated by * (P < 0.05).

**Figure 8.**
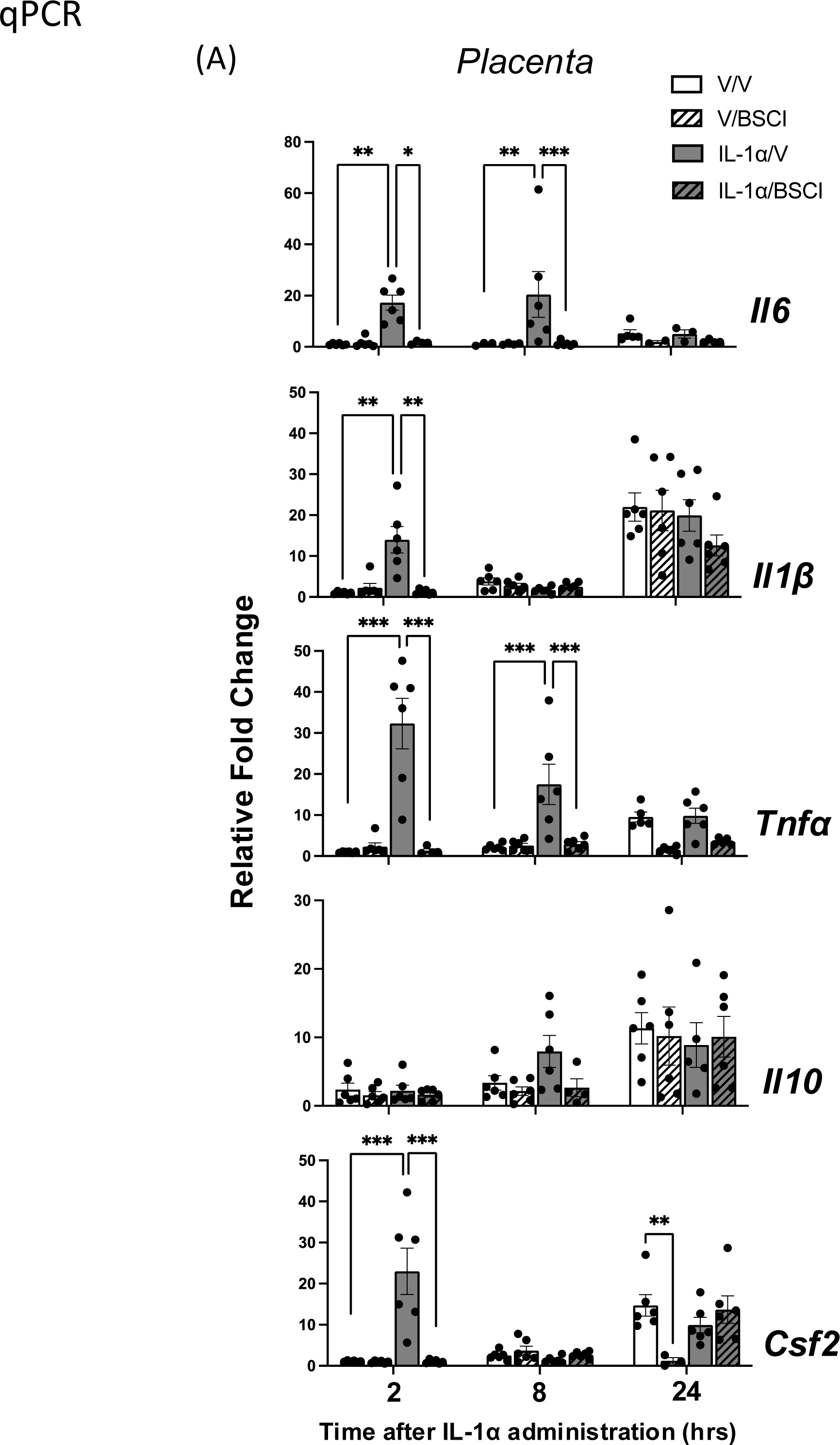
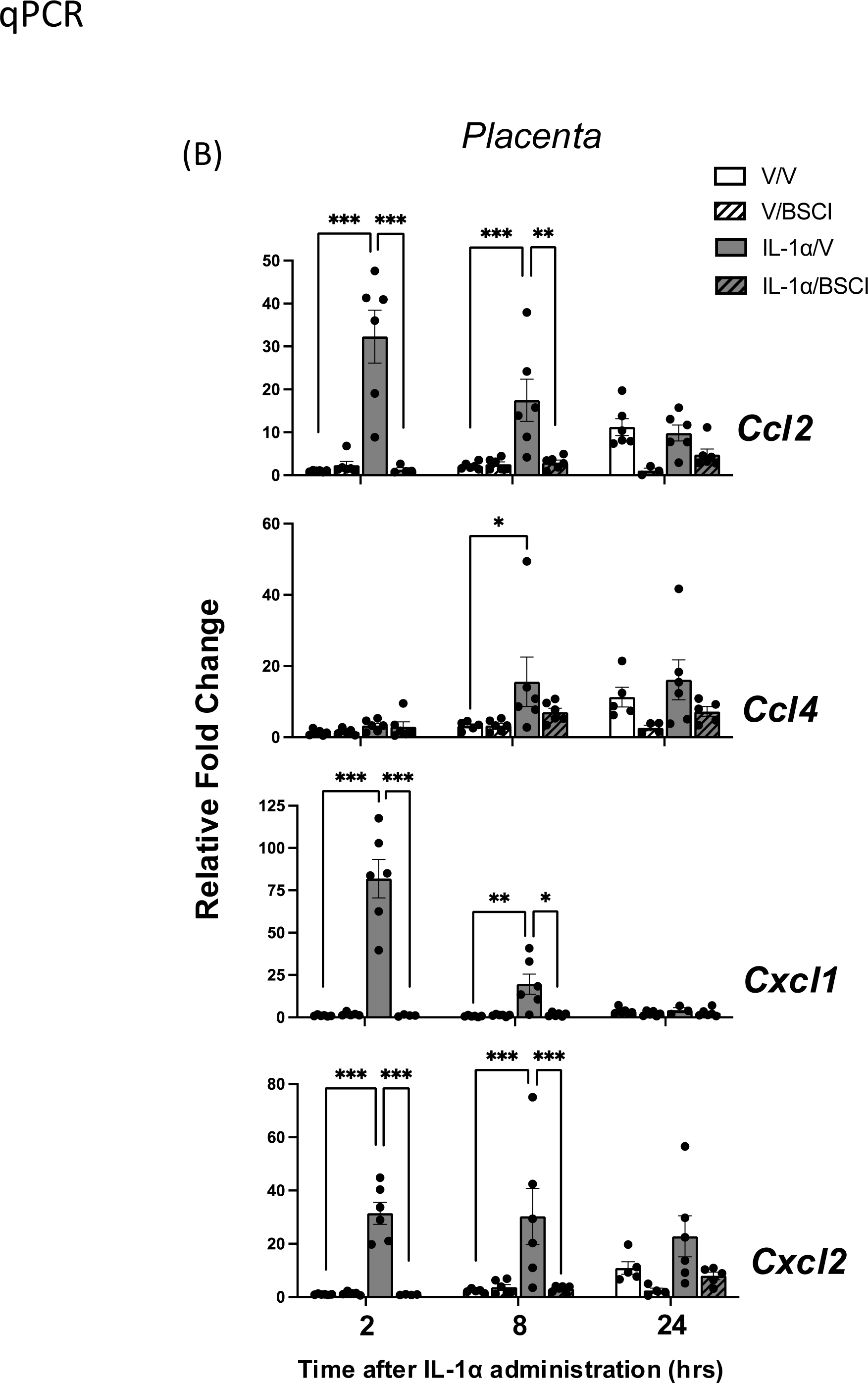
Changes in cytokine mRNA levels in the mouse placenta of GD16 and GD17 pregnant mice following local IL-1α intra-amniotic administration and pretreatment with Broad Spectrum Chemokine Inhibitor (BSCI). **(A)** Transcript levels of pro-inflammatory (*Il6*, *Il1*β, *Tnf*α, *Csf2/Gmscf*) and anti-inflammatory (*Il10*) cytokines; **(B)** Chemokines (*Ccl2/Mcp1*, *Ccl4/Mip1*β, *Cxcl1/KC/Gro*α, and *Cxcl2/Mip2*α) were detected by Real-Time RT-qPCR. Shown are vehicle (white bars), BSCI-treated (striped bars), IL-1α-injected (grey bars), and IL-1α-injected BSCI-treated samples (striped grey bars), n = 5–6/group. Two-way ANOVA was utilized, followed by a Bonferroni post-test. Results were expressed as mean ± SEM. Significant difference is indicated by * (P < 0.05), ** (P < 0.01), and *** (P < 0.001).

### Genomic and Transcriptomic Profiling Shows BSCI Prevents IL-1α-Driven Inflammatory Reprogramming in the Pregnant Myometrium

To determine if the changes in inflammatory gene expression, and subsequent suppression, occurred at a genome-wide scale, we first performed RNA sequencing (RNA-seq) on myometrial tissues collected 2 hr after intra-amniotic injection for Vehicle, BSCI, IL-1α, IL-1α+BSCI groups. Genes examined by RT-qPCR followed similar trends by RNA-seq. For example, increased gene expression in response to IL-1α was exhibited in *Ptgs2* (fold change = 9.78, padj = 1.15×10^-5^), *Il1b* (fold change = 30.48, padj = 1.33×10^-4^), and *Nfkb1* (fold change = 4.98, padj = 8.97×10^-4^), and this was suppressed by pretreatment with BSCI in the IL-1α+BSCI group (**Fig. 9A-C, Table S1-2**). No significant change in expression for these genes was exhibited in BSCI alone compared to vehicle (**Fig. 9A-C, Table S3**). Genome-wide, differential expression analysis confirmed this pattern, compared with vehicle treatment, IL-1α induced significant differential expression of 911 genes at 2 hr (abs. fold change ≥ 2, padj < 0.05) (**Fig. 9D, Table S1**), IL-1α+BSCI blunted this response with 341 differentially expressed genes (**Fig. 9E, Table S2**) while just two differentially expressed genes were identified in BSCI alone, one increasing expression (predicted, non-protein coding *Gm6361*) and one decreasing expression (predicted gene *Gm14440*) compared to vehicle (**Fig 9F, Table S3, Fig. S1A**). Of the 911 genes differentially expressed at 2 hrs post-IL-1α treatment, 559 genes were upregulated, including the inflammatory and pro-labor transcripts examined by RT-qPCR above (*Ccl2, Ccl4, Cxcl2, Il6, Il1b, Tnf, Csf2, Nfkb1, Ptgs2)* and 352 downregulated (abs. fold change ≥ 2, padj < 0.05). In line with this, Gene Ontology (GO) analyses of genes increasing expression in response to IL-1α had enrichment of inflammation-linked terms, including cytokine activity, tumor necrosis factor receptor binding and activation of immune cells (lymphocyte activation, T cell activation, and T cell differentiation) (**Table S4**). In the 352 genes downregulated at 2 hr in response to IL-1α, GO terms were primarily linked to transcriptional (transcription by RNA polymerases II, DNA binding) and metabolic processes (regulation of primary metabolic process) (**Table S5**).

**Figure 9.**
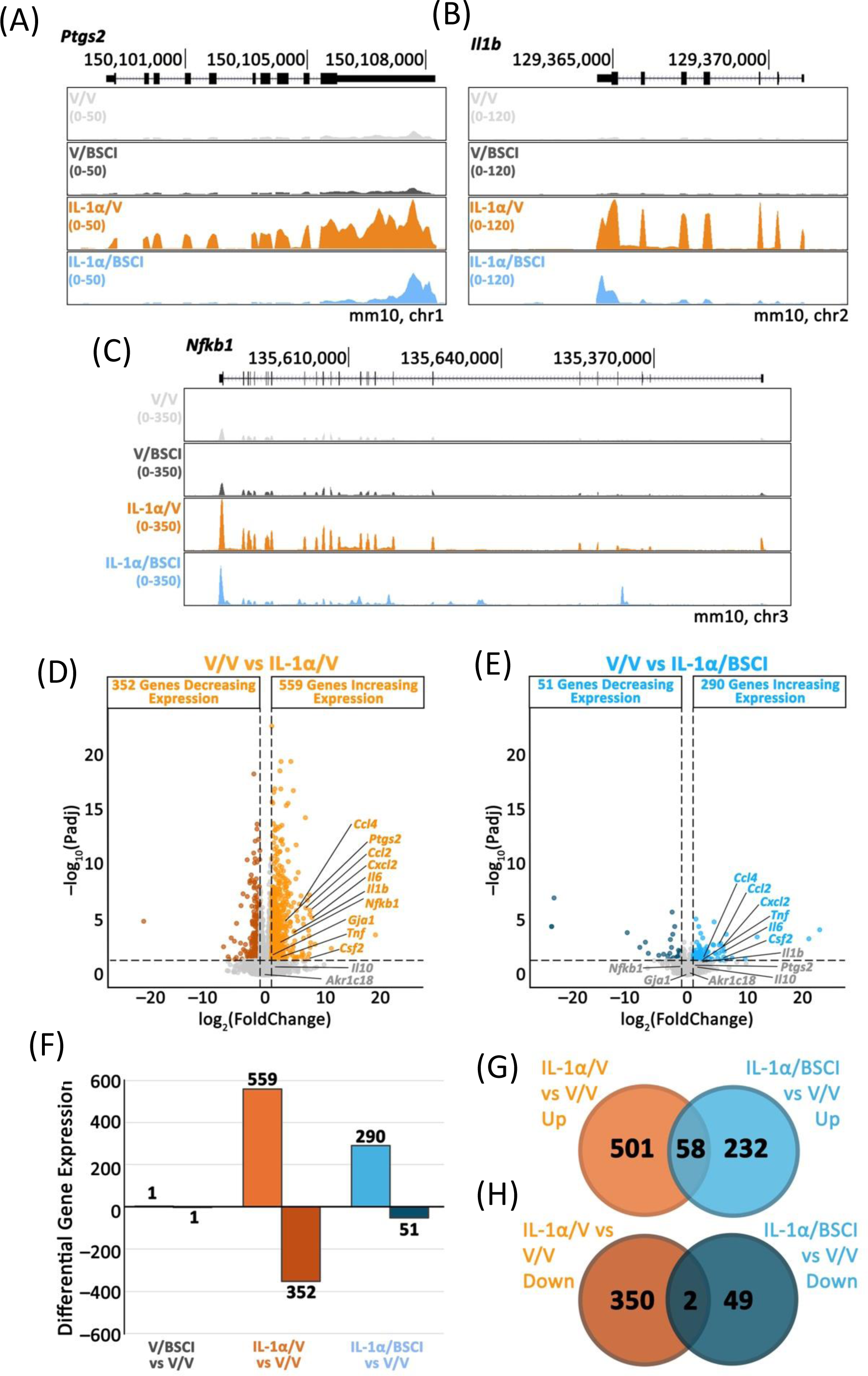
Broad Spectrum Chemokine Inhibitor (BSCI) blunts IL-1α-driven changes in transcriptomic response in the pregnant myometrium at 2 hr. RNA-seq was performed on myometrial tissue collected 2 hr after intra-amniotic injection of vehicle (saline) or IL-1α (400 ng/sac), with or without daily BSCI pretreatment (10 mg/kg). **(A-C)** UCSC bedgraph tracks of transcript reads mapped to the mm10 genome for representative pro-labor and inflammatory genes *Ptgs2*, *Il1b*, and *Nfkb1*. **(D)** Volcano plot of differentially expressed genes for V/V vs IL-1α/V (559 genes upregulated, 352 downregulated; abs. fold change ≥ 2, padj < 0.05). Significance thresholds are represented by dashed lines on graphs, and pro-labor and inflammatory genes examined by RT-qPCR are annotated. **(E)** Volcano plot of differentially expressed genes for V/V vs IL-1α/BSCI (290 upregulated, 51 downregulated). Pro-labor and inflammatory genes examined by RT-qPCR are annotated). **(F)** Bar graph summarizing the number of differentially expressed genes across treatment comparisons (V/V vs V/BSCI, V/V vs IL-1α/V, V/V vs IL-1α/BSCI). **(G)**Venn diagrams showing the overlap of upregulated and **(H)** downregulated genes between the V/V vs IL-1α/V and V/V vs IL-1α/BSCI comparisons. n = 3 biological replicates per group.

Pretreatment with BSCI in IL-1α-challenged animals substantially blunted the transcriptomic response. The number of genes increasing expression dropped to 290 (abs. fold change ≥ 2, padj < 0.05). GO terms enriched in these genes still included those associated with inflammatory response (e.g. cytokine activity, chemokine activity), but no longer exhibited terms linked to immune cell processes (**Table S6**), demonstrating that BSCI selectively suppresses the IL-1α-driven inflammatory transcriptional program at 2 hrs. In the 51 down-regulated genes in the IL-1α + BSCI group, GO terms for ion binding, small molecule binding and collagen binding were enriched (**Table S7**). Analyses of upregulated genes showed significant overlap; 58 of the 559 IL-1α-induced upregulated genes were shared with the IL-1α/BSCI group (Representation factor (RF)=8.1, P = 7.86×10^-36^; **Fig. 9G**), whereas just 2 of 352 downregulated genes (RF=2.5, P=0.19; **Fig. 9H)**. Of the 58 upregulated gene that are shared between IL-1α and IL-1α/BSCI the magnitude of increased expression was reduced in IL-1α/BSCI for 50 of these genes, indicating BSCI is able to suppress genome-wide changes in IL-1α-induced changes in gene expression.

Given that BSCI appeared to suppress genome-wide IL-1α−induced inflammatory-associated gene expression changes in the pregnant myometrium, we next examined if this effect was sustained at 24 hrs. By 24 hrs, the majority of genes that exhibited upregulation by IL-1α were suppressed by BSCI. For example, matrix metalloproteinase 3 (*Mmp3*), previously shown to be upregulated in mouse, rat, and human myometrium at labour onset (78–80), was elevated in response to IL-1α at 24 hrs (fold change = 12.13, padj = 2×10^-20^), but this elevation was suppressed in IL-1α+BSCI myometrium and no longer significantly different from vehicle (fold change = −0.90, padj = 0.646; **Fig. 10A**,).

**Figure 10.**
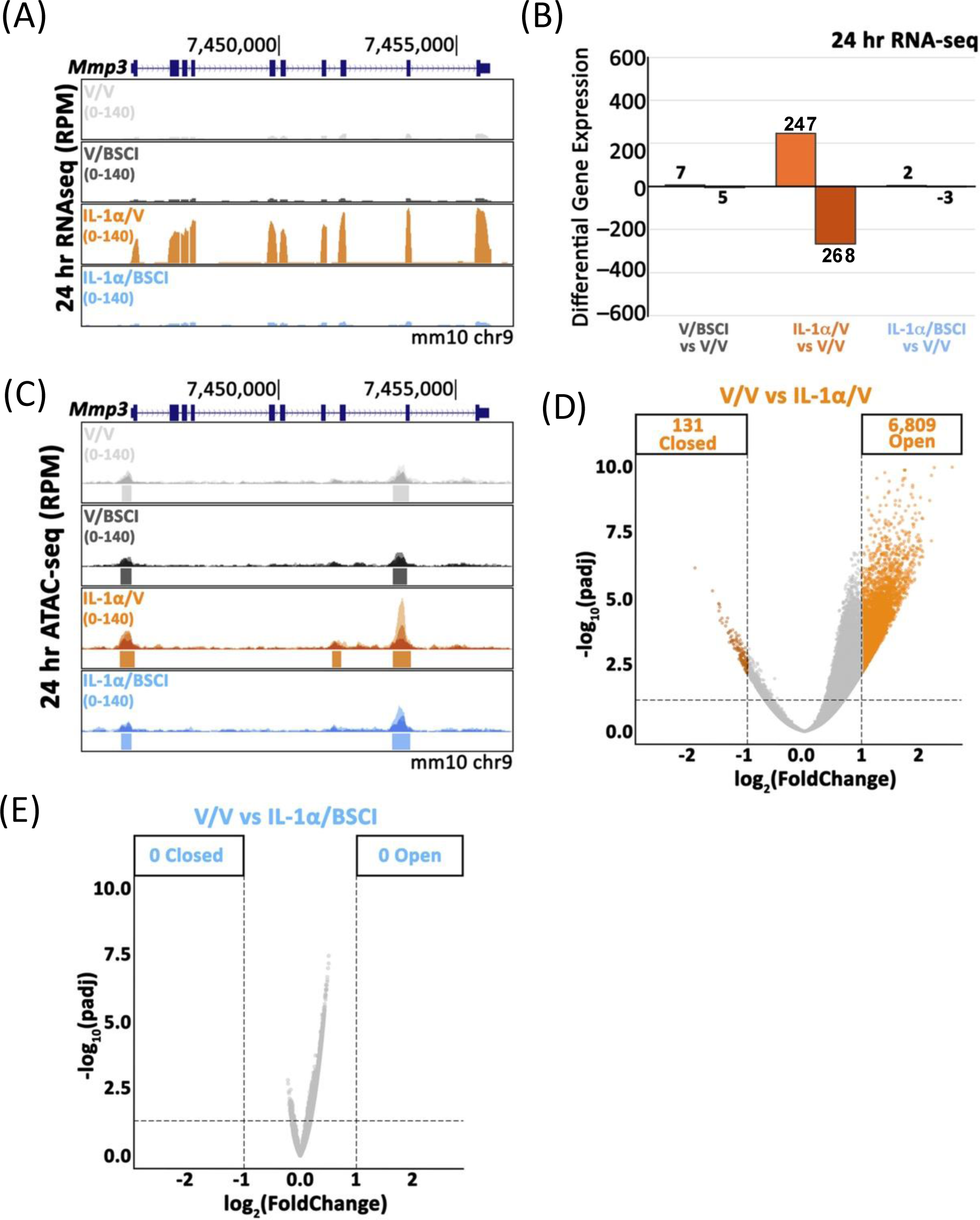
Broad Spectrum Chemokine Inhibitor (BSCI) blocks IL-1α-driven changes in the transcriptome and associated chromatin accessibility in the pregnant myometrium at 24 hrs. RNA-seq and ATAC-seq were performed on myometrial tissue collected 24 hr after intra-amniotic injection of Vehicle (saline) or IL-1α (400 ng/sac), with or without daily BSCI pretreatment (10 mg/kg). **(A)** UCSC bedgraph track of 24-hr RNA-seq reads (RPM, mm10 genome) at the *Mmp3* locus showing fewer reads in V/V and V/BSCI, increased reads aligning with *Mmp3* in IL-1α/V and again fewer reads with BSCI pretreatment (IL-1α/BSCI). **(B)** Bar graph summarizing differentially expressed genes from 24-hr RNA-seq (abs. fold change ≥ 2; padj < 0.05) across V/V vs V/BSCI, V/V vs IL-1α/V, and V/V vs IL-1α/BSCI comparisons; the number of differentially expressed genes is listed on the Y-axis and is shown above each bar. **(C)** UCSC bedgraph track of 24-hr ATAC-seq reads (RPM, mm10 genome) at the *Mmp3* locus showing increased chromatin accessibility showing fewer reads in V/V and V/BSCI, increased reads aligning with *Mmp3* in IL-1α/V and again fewer reads with BSCI pretreatment (IL-1α/BSCI).. Bars under tracks indicate MACS2-called peaks. **(D)** Volcano plot of differentially accessible chromatin regions for V/V vs IL-1α/V (131 closed, 6,809 open; abs. fold change ≥ 2, padj < 0.05). **(E)** Volcano plot of differentially accessible chromatin regions for V/V vs IL-1α/BSCI (0 closed, 0 open). Y-axis is -log_10_(padj) and X-axis log_2_(foldchange); dashed lines represent significance thresholds. Samples in grey did not meet criteria for significance. n = 3 biological replicates per group.

In line with this, whereas 515 genes were differentially expressed 24 hrs after IL-1α treatment, in IL-1α+BSCI myometrium only 5 genes displayed differential expression at 24 hrs (abs. fold change ≥ 2, padj < 0.05) compared to vehicle controls (**Fig. 10B, S1C-D, Table S8, 9**). BSCI alone compared to vehicle displayed 12 differentially expressed genes (**Fig. S1B, Table S10**). Of the 515 genes differentially expressed 24 hrs after IL-1α treatment, 247 genes showed significantly increased expression and GO enrichment demonstrated the association of these upregulated genes with cytokine and chemokine activity (e.g., cytokine activity, cytokine receptor activity, cytokine receptor binding, chemokine-mediated signaling pathway), metalloendopeptidase activity, and prostaglandin receptor activity (**Table S11**). GO enrichment of the 268 genes with significantly decreased expression in response to IL-1α, included calcium ion binding (e.g., calcium ion binding involved in regulation of cytosolic calcium ion concentration) and cell-cell signaling (**Table S12**), likely involved in modulating smooth muscle contractility. Considering the 5 genes with altered expression in IL-1α+BSCI compared to Vehicle, two genes with increased expression, (*Gm6565* is non-coding, and *Gm14308* is a predicted gene), and three genes significantly decreased expression (*Gm43951*, predicted gene *Gm2026,* and *Epha4*), none have known roles in labour onset. Due to the small number of genes changing expression, no GO enrichment was present in IL-1α+BSCI vs Vehicle myometrium, as is also the case for BSCI vs Vehicle. These data demonstrate that IL-1α induces an inflammatory response at the transcriptomic level linked with labor onset that is completely blocked by pretreatment with BSCI at 24 hrs, while BSCI vehicle control animals exhibited minimal transcriptomic changes in the pregnant myometrium.

We were interested to determine if changes to accessible chromatin occurred in the myometrium after IL-1α treatment and if these were blunted in IL-1α+BSCI treatment. We conducted an assay for transposase-accessible chromatin with sequencing (ATAC-seq). As with RNA-seq, large-scale changes in chromatin accessibility were identified after IL-1α injection that were not present in the IL-1α+BSCI myometrium. For example, *Mmp3* chromatin was less accessible in vehicle control and BSCI groups, while a prominent peak region in intron eight gained accessibility in response to IL-1α (**Fig. 10C**). In IL-1α+BSCI this increased accessibility was not observed at intron eight (**Fig. 10C**). No significant changes in chromatin accessibility occurred in BSCI vs vehicle groups (**Table S13**) while a total of 6,940 chromatin regions were significantly altered in response to IL-1α (abs. fold change ≥ 2, FDR < 0.05); 6,809 increasing accessibility and 131 chromatin regions losing accessibility compared to vehicle controls (**Fig. 10D, Table S14**). Strikingly, IL-1α+BSCI groups exhibited no significant changes in chromatin accessibility compared to vehicle controls (**Fig. 10E, Table S15**). These data further support the ability of BSCI to suppress gene expression and imply that this may, in part, be due to the prevention of changes in chromatin accessibility. In summary, BSCI pretreatment prevented the widespread changes in chromatin accessibility and gene expression induced by IL-1α in the pregnant mouse myometrium, while having no effect on these parameters on its own.

### IL-1α-induced Changes in Uterine Resident Macrophages

We examined the presence and polarization status of macrophages in the pregnant mouse uterus collected 24 hrs after intra-amniotic IL-1α infusion using immunohistology. Immunostaining was performed using antibodies for F4/80, a 160 kDa plasma membrane glycoprotein and marker of murine macrophages (81). Using QuPath imaging software, we found that in IL-1α-challenged mice, there was a significant increase in the number of F4/80+ cells in the mouse myometrium (3-fold increase) and decidua (2-fold increase) as compared to vehicle (saline) treated animals. Notably, the number of macrophages in myometrium (avg. 9400 cells per mm^2^) was nearly twice as high as in decidua (avg. 5424 per mm^2^) (P < 0.01-0.001, **Fig. 11**). We found that the BSCI pretreatment of pregnant mice significantly decreased the total number of F4/80+ cells only in the IL-1α-inflamed myometrium (P < 0.01, **Fig. 11**), but not in the decidua, and did not affect the immune cell detection in control Vehicle-treated pregnant murine uterus.

**Figure 11.**
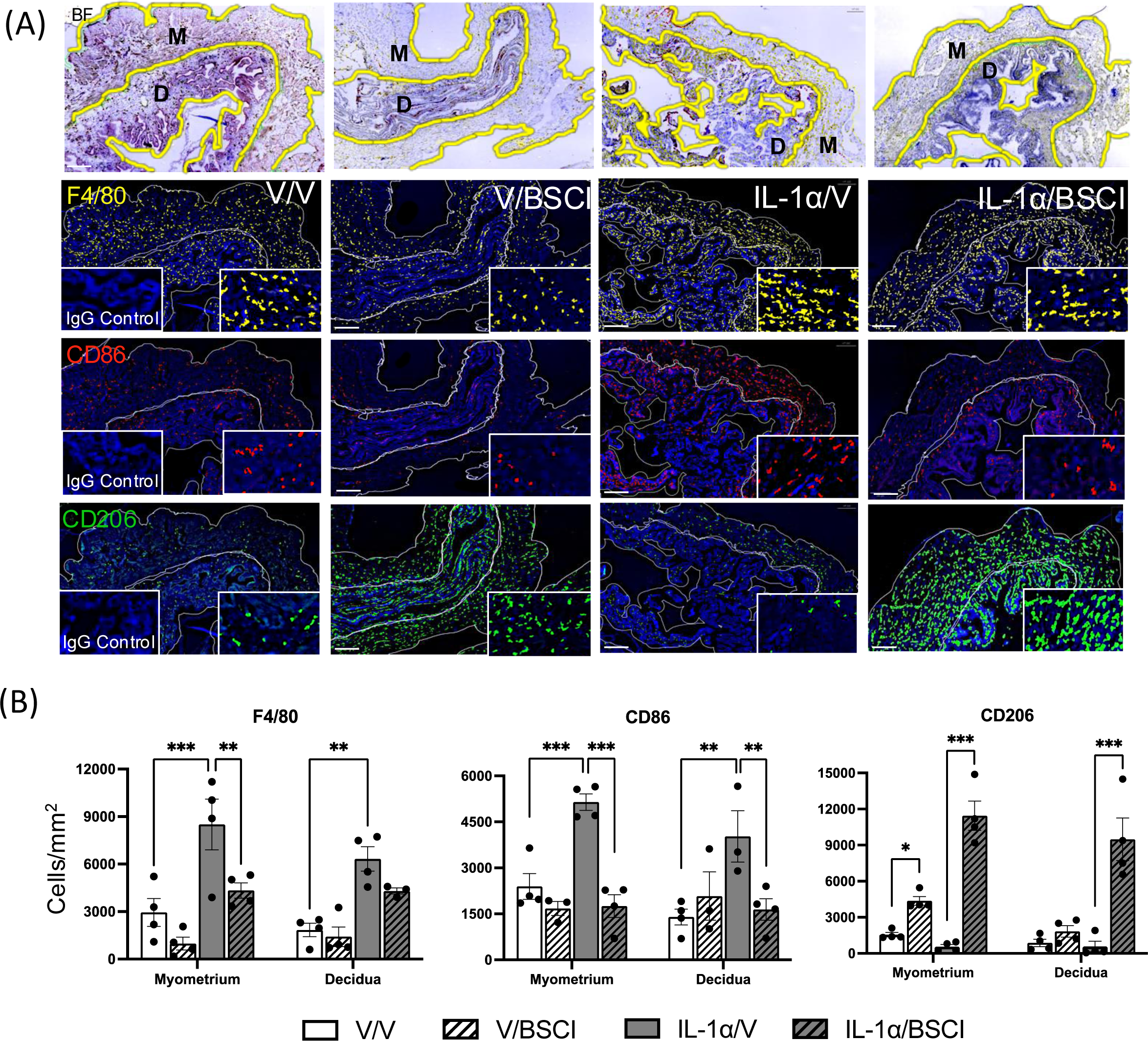
Monocyte infiltration and macrophage polarization in the mouse uterus 24 hrs following local IL-1α intra-amniotic administration and pretreatment with Broad Spectrum Chemokine Inhibitor (BSCI). Macrophages were identified using anti-F4/80 antibody (yellow), using anti-CD86 for pro-inflammatory M1 macrophages (red), and using anti-CD206 for homeostatic M2 macrophages (green). Bright field (BF, top row) images were used to mask the borders between the myometrium and the decidua. **(A)** Shown are the representative immunofluorescence images with negative control inlets (images taken with respective IgG controls for each antibody). **(B)** Qupath software was used to quantify the number of macrophages in the myometrium and decidua. The bar graphs show quantification of vehicle (white bars), BSCI-treated (striped bars), IL-1α-injected (grey bars), and IL-1α-injected BSCI-treated samples (striped grey bars), n = 5–6/group. Two-way ANOVA was utilized, followed by a Bonferroni post-test. Results were expressed as mean ± SEM. Significant difference is indicated by ** (P < 0.01) and *** (P < 0.001). Nuclei were counterstained with DAPI (blue). The scale bar represents XX μm.

To quantify the polarization status of uterine macrophages by immunohistology, we used two specific markers: (1) CD86 for pro-inflammatory M1-like macrophages and (2) CD206 for homeostatic M2-like macrophages. Not surprisingly, we found that the number of CD86+ cells was significantly (P < 0.01-0.001) increased in both uterine compartments, myometrium (2.2-fold) and decidua (2.9-fold) of laboring IL-1α-injected mice compared to control vehicle-injected non-laboring animals. Importantly, we detected that the number of CD86+ M1-like macrophages was not elevated in the uterine tissues of IL-1α+BSCI treated mice and was similar to M1-macrophage number in control Vehicle-treated animals (P < 0.01-0.001, **Fig. 11**). Moreover, we detected that the number of CD206+ M2-like macrophages significantly increased in the myometrium (7-fold) and decidua (10-fold) of the IL-1α+BSCI treated compared to control Vehicle-treated mice (P < 0.001, **Fig. 11**). In the BSCI control group, we found a significant increase in myometrial CD206+ macrophages (2.8-fold, P < 0.05, **Fig. 11**). Together, these findings show that IL-1α increased uterine macrophage abundance and drove polarization toward the pro-inflammatory M1-like phenotype. BSCI pretreatment lowered macrophage numbers in the inflamed myometrium and favoured the homeostatic M2-like phenotype.

## DISCUSSION

Our study determined that the BSCI compound FX125L can partially reduce the incidence of PTB in a mouse model of sterile IAI induced by the injection of the alarmin, IL-1α. IL-1α was the first cytokine linked to both term and preterm labor (15, 17, 19, 26); it is constitutively expressed in different cell types (epithelial, stromal and muscle cells, as well as leukocytes) and is released upon membrane damage, where it functions as both a transcriptional regulator and an extracellular signal. Once released, IL-1α engages Toll-like and NOD-like pathways to recruit neutrophils and monocytes and to initiate a self-amplifying cytokine and chemokine cascade (15, 25). Elevated IL-1α levels are reported in “sterile” inflammatory conditions such as rheumatoid arthritis and ischemia-reperfusion injury and have been linked to term and preterm labor initiation (15, 82). Sterile IAI accounts for nearly one-third of sPTB with intact membranes and is characterized by elevated levels of inflammatory mediators and the absence of detectable microbes (3, 6, 20).

Early animal studies used systemic IL-1α injections to induce sPTB and later shifted to the intra-uterine route to confine the model to the local uterine inflammation, closely mimicking human pathology (24, 27, 28). Ultrasound-guided intra-amniotic injection of IL-1α was recently used to create a pathophysiologic PTB model with an inflammation localized inside the amniotic sac while minimizing systemic spillover (16). This approach mirrors the clinical pattern in pregnant women, where inflammatory mediator levels rise first in amniotic fluid, fetal membranes, decidua, and myometrium, with maternal plasma changes that can be modest or delayed (23, 26). It also standardizes dose per amniotic sac and supports controlled assessment of early tissue responses before labor, which helps separate the cause from the consequence of the labor process. Our data confirm that intra-amniotic administration of IL-1α (400 ng/amniotic sac) at GD16.5 in C57BL/6 mice reliably induces PTB within approximately 24 hrs and recreates the coordinated inflammatory response of sterile IAI in humans. Within 2 hrs, we observed rapid rises in pro-inflammatory cytokines and chemokines across maternal plasma, myometrium, decidua, placenta, and amniotic fluid, including IL-1α, IL-1β, IL-6, IL-12p40, IL-12p70, TNFα, CSF2, CSF3, CCL2, CCL4, CXCL1, CXCL2, and CCL11, along with anti-inflammatory IL-10. Myometrium and decidua showed temporally distinct profiles: in the myometrium IL-1α induced a transient increase in inflammatory gene and protein expression with levels returning to baseline by 8 hrs after which there was an increase in the expression of genes critical for myometrial contraction (e.g. the gap junction protein, *Gja1*) (83–85). In contrast, the induction of inflammatory gene and protein expression in the decidua was more sustained, with many expressed at high levels through to preterm labor (79–82). This sequence supports compartmental roles for these uterine tissues. In the myometrium, inflammation induces proteins that prepare the muscle for contraction (“activation”), while in the decidua, sustained inflammation results in the production of uterotonic agonists that “stimulate” myometrial contractions (83–85). Using quantitative immunofluorescence imaging, we also found marked increases in F4/80+ macrophages in both the myometrium and the decidua, consistent with acute leukocyte recruitment and local activation driving these processes.

Importantly, treatment with the BSCI, FX125L prevented sterile PTB in 64% of animals. We selected BSCI because it prevented infection-mediated PTB in mice and in non-human primate models (46, 47). In the current study, pretreatment reduced preterm delivery, extended latency till term, and in half of the cases preserved neonatal viability in dams that delivered at term. These outcomes mirror our earlier LPS mouse study, which showed that BSCI reduced cytokines in maternal tissues and the amniotic compartment and prevented PTB in 64% of cases (46). LPS (i.e., endotoxin), delivered intraperitoneally, activates TLR4 and induces an early systemic cytokine response across maternal compartments with secondary uterine and placental effects (46). Here, intra-amniotic IL-1α creates a steep local gradient within the amniotic cavity and fetal membranes, yielding early peaks in myometrium and decidua with modest spillover into plasma and liver, a profile that better reflects sterile IAI.

Protection from IAI and PTB induction by BSCI was partial. Dams either delivered preterm within 24 hrs or carried their litters past GD18. Nine of 14 dams that preventively received BSCI and were induced with IL-1α carried pregnancy to term; live pups were recorded in 7 of 9 litters. Five of 14 BSCI-pretreated dams delivered preterm 18 to 24 hrs after IL-1α injection, within the same window as the vehicle-treated IL-1α group. Pups delivered prematurely were without vital signs. Fetal demise was also noticed in the “BSCI rescued” group, where 2 of the 9 term litters had no pups with vital signs, and most pups in those litters were undeveloped. Therefore, uterine quiescence and fetal wellbeing were two separable outcomes in this mouse model. BSCI was able to keep the uterus quiescent in most dams, while fetal demise still occurred in a subset of pregnancies subjected to the IAI with IL-1α injection.

This incomplete rescue is informative in two respects. First, it highlights the strength of the inflammatory stimulus. A single 400 ng dose of IL-1α per amniotic sac induced delivery within approximately 24 hrs in all vehicle-treated dams, with no viable pups in any litter. The same challenge overcame daily systemic BSCI pretreatment in approximately one-third of the animals, resulting in fetal demise. These findings underscore the intensity of the inflammatory challenge and suggest that, by the time labor was initiated, the inflammatory cascade had advanced substantially. Second, the results demonstrate partial protection achieved by BSCI. The remaining BSCI-pretreated dams maintained their pregnancies to term despite the maximal single-alarmin challenge. This provides a proof of principle that BSCI blockade can counteract the inflammatory mechanism that would otherwise result in uniformly fatal fetal outcome in this model. However, the BSCI dose and treatment timing were fixed in this study design and were not optimized relative to the IL-1α challenge. Whether a higher BSCI dose or earlier initiation of anti-inflammatory treatment can further improve the PTB rate remains to be determined.

Fetal and placental weights provide additional insight into the extent of rescue. Pup weights in IL-1α/BSCI dams that carried their pregnancies to term were generally comparable to the vehicle and BSCI-only groups. Daily BSCI treatment alone also had no effect on litter weight (**Table 1**), suggesting that BSCI does not adversely affect fetal growth under the conditions tested. In contrast, litter weights in the preterm IL-1α/BSCI pregnancies were below those of litters with live-born pups, consistent with delivery at an earlier gestational age (**Fig. 2D**). Placental weight comparisons are complicated by differences in gestational age at collection, as preterm placentas were collected on GD17, whereas term placentas collected between GD18.5-20 (**Table 1**).

Within the IL-1α/BSCI group, placental weights varied considerably. Notably, the single abnormally large placenta (0.68 g) was from a term litter in which no pups showed vital signs, raising the possibility that placental pathology contributed to intrauterine fetal demise following IL-1α-induced IAI. However, placental histology was not performed in this study, and the underlying pathology remains unknown. Systematic histological comparison of placentas from rescued and non-rescued pregnancies will be necessary to determine whether placental abnormalities are associated with failed rescue and to better define the limits of BSCI-mediated protection.

Our earlier non-human primate results strengthen this conclusion by showing coordinated suppression of infection-induced inflammation in both maternal and fetal systems (47). In that study, BSCI lowered amniotic fluid cytokines IL-6 and IL-8, decreased cytokine levels in fetal tissues, and reduced maternal inflammatory markers while maintaining uterine quiescence during an active infectious challenge (47). This result matches what we observe here in a sterile mouse PTB model, where BSCI decreased IL1α-induced cytokine and chemokine induction in plasma, myometrium, decidua, placenta, and amniotic fluid, in particular, amniotic IL-6 levels. Importantly, in pregnant women, IL-6 levels in amniotic fluid are a clinical benchmark for detecting IAI associated with imminent PTB (86, 87). Thus, decreases in maternal and fetal inflammatory markers and preservation of fetal growth support a key takeaway from our study: BSCI limits leukocyte infiltration and inflammation within the fetoplacental unit, stabilizes the intra-amniotic environment, and maintains uterine quiescence even in the presence of strong danger signals.

BSCI reduced myometrial activation from 2 hrs post-injection until full term. After intra-amniotic injection of IL-1α, *Nfkb1, Ptgs2, Akr1c18*, and *Gja1* all showed strong induction in the myometrium. The gene-specific pattern aligns with known mechanisms of labor initiation (84, 88). BSCI pretreatment significantly reduced all transcripts tested by RT-qPCR at 2 hrs, interrupting the acute inflammatory signaling. Lowering *Nfkb1* limits transcriptional increase in inflammatory genes; suppressing *Ptgs2* reduces prostaglandin synthesis capacity; decreasing *Akr1c18* preserves progesterone signaling by limiting 20α-HSD activity (89); reducing *Gja1* constrains Connexin-43 availability for electrical coupling. Together, these data show that BSCI blocks the early transcriptional cascade that links inflammation to prostaglandin production, progesterone withdrawal pathways, and gap-junction assembly.

RNA-seq and ATAC-seq data show that intra-amniotic IL-1α drives a broad shift in transcription that is accompanied by widespread gains in chromatin accessibility, consistent with prior evidence that labor-relevant myometrial programs are regulated through transcriptional remodeling and mechano-endocrine integration (90–92). This analysis linked inflammatory signalling to gene regulation in the pregnant myometrium. 2 hr and 24 hr RNA-seq data show that intra-amniotic IL-1α drives a large-scale shift in transcription, with strong upregulation of cytokines, chemokines, and contraction-associated genes, in line with the RT-qPCR and protein findings. BSCI largely suppressed these IL-1α induced changes in gene expression at 2 hrs, with virtually no changes in the transcriptome in IL-1α+BSCI at 24 hrs. BSCI also suppressed changes in chromatin accessibility induced by IL-1α at 24hrs.

We further examined the potential of the BSCI to influence local uterine macrophage populations, which have been linked to labor onset (56, 93). Studies in humans and mice report that homeostatic M2-like macrophages, identified by CD206 expression, are enriched during late gestation and decline with the onset of labor (58–60, 64). Conversely, pro-inflammatory M1-like macrophages, marked by CD86, increase during parturition (52, 55, 57). Our results show that in the IL-1α-induced pregnant mice, the number of F4/80^+^ CD86^+^ macrophages rose in myometrium and decidua, while BSCI reduced macrophage accumulation and returned the CD86 signal to control levels. The most notable change was the expansion of CD206^+^ cells in myometrium and decidua in BSCI-treated animals, including the IL-1α/BSCI group. This pattern aligns with our prior *in vitro* findings that media conditioned by BSCI-treated myocytes polarized human macrophages toward an M2-like phenotype (51). The combination of reduced influx of F4/80^+^ CD86^+^ cells and an increase in F4/80^+^CD206^+^ cells suggests that BSCI promotes the resolution of inflammation. Complementing these observations, our recent study identified uterine M1 macrophages, but not M2 cells, as a local source of IL-1α that directly influences myometrial contractility (94).

Although the present study highlights the potential of BSCIs as a promising novel class of therapeutics to prevent inflammation-induced preterm labor in patients with sterile IAI and intact membranes, it has limitations. The intra-amniotic IL-1α mouse model is clinically relevant, yet human sPTB often reflects overlapping processes that evolve over weeks, including parallel rises in other alarmins, progressive cervical remodeling, and placental vascular or endocrine changes (12, 95). A single alarmin at a fixed gestational day cannot capture that diversity, and procedural variables such as the number of sacs injected, anesthesia, and handling may contribute to low-grade inflammation that interacts with IL-1α. In addition, our immune phenotyping centred on macrophages, using F4/80, CD86, and CD206 to study dominant M1/M2 states, which is a practical but simplified approach. Macrophage phenotypes are plastic and span a continuum that shifts with local cues, making state assignments reversible and context dependent (96). Chemokine tone can reprogramme these cells within hrs, so broader panels and temporal profiling are needed to capture transitions relevant to labor. Future *in vivo* studies, including single-cell RNA sequencing with spatial mapping and lineage tracing, would resolve cell identities, origins, and trajectories and place BSCI effects within the broader cellular network. Finally, translational progress requires pharmacokinetics, placental transfer, and biodistribution data for BSCI in late gestation, as well as assessment of maternal reproductive health and offspring outcomes beyond birth, including growth, neurodevelopment, and sexual maturation.

## Author Contributions

Conceptualization and project design: O.S. and S.L.; methodology: A.B-R and A.D.; investigation: A.B.-R., N.B., I.C., and A.D.; formal analysis: A.B.-R, N.B., and Z. E. G.; writing: A.B.-R., O.S., Z.E.G., and J.A.M; writing-review and editing: D.F., D.G., O.S., J.A.M., and S.L.; supervision: S.L. and O.S. All authors have read and agreed to the published version of the manuscript.

## Data Availability Statement

The original contributions presented in this study are included in the article/Supplementary Material. Further inquiries can be directed to the corresponding author(s). RNA-seq and ATAC-seq data have been deposited in the Gene Expression Omnibus (GEO) under accession numbers GSE317954 (RNA-seq) and GSE292301 (ATAC-seq).

## Acknowledgments

We want to thank Louise Brown of the Network Biology Collaborative Centre Advanced Imaging Facility at the Lunenfeld-Tanenbaum Research Institute. We thank David Fox (Warwick University, UK) for the generous gift of the BSCI (FX125L).

Conflicts of Interest

The authors declare no conflicts of interest.

---

**Supplemental Figure 1:**
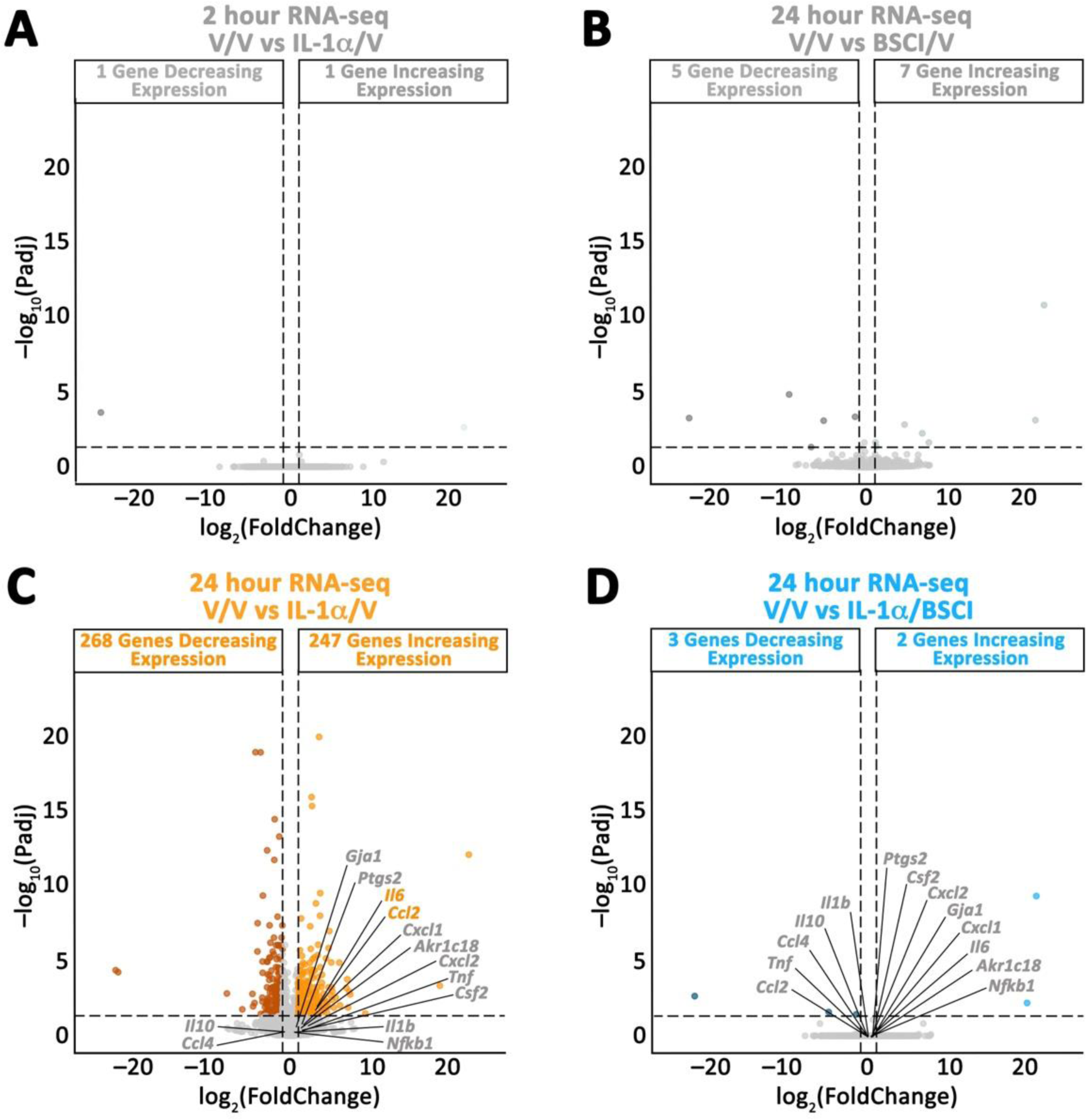
BSCI suppresses genome-wide transcriptomic changes induced by IL-1α at 2 hrs and 24 hrs post-intraamniotic injection in the pregnant mouses myometrium. RNA-seq was performed on myometrial tissue collected 2 hr and 24 hr after intra-amniotic injection of vehicle (saline) or IL-1α (400 ng/sac), with or without daily BSCI pretreatment (10 mg/kg). **(A)** Volcano plot for V/V vs BSCI/V demonstrating 1 gene increasing and 1 gene decreasing expression at 2 hrs (abs. fold change ≥ 2, padj < 0.05). Volcano plots for gene expression at 24hrs in **(B)** V/V vs BSCI/V, **(C)** V/V vs IL-1α/V, and **(D)** V/V vs IL-1α/BSCI comparisons. Genes examined by RT-qPCR are called out in **(C)** and (**D).** Thresholds for significance are represented by dashed lines, where abs. fold change ≥ 2 and padj < 0.05.

